# Mitochondrial calcium uniporter regulates human fibroblast-like synoviocytes invasion via altering mitochondrial dynamics and dictates rheumatoid arthritis pathogenesis

**DOI:** 10.1101/2024.08.13.607857

**Authors:** Lakra Promila, Kabita Sarkar, Shivika Guleria, Manisha Rathore, Nishakumari C Singh, Shazia Khan, Manendra Singh Tomar, Veena Ammanathan, Manoj Kumar Barthwal, Jagavelu Kumaravelu, Ashutosh Shrivastava, Kalyan Mitra, Rajdeep Guha, Amita Aggarwal, Amit Lahiri

**Affiliations:** Pharmacology Division, CSIR-Central Drug Research Institute, Lucknow, 226031, India; Academy of Scientific and Innovative Research (AcSIR), Ghaziabad-201002, India; Sophisticated Analytical Instrument Facility and Research Division, CSIR-Central Drug Research Institute, Lucknow, India; Lab Animal facility, CSIR-Central Drug Research Institute, Lucknow, India; Department of Clinical Immunology and Rheumatology, Sanjay Gandhi Postgraduate Institute of Medical Sciences, Lucknow, India; Centre for Advance Research, Faculty of Medicine, King George Medical University, Lucknow, India

**Keywords:** Rheumatoid arthritis, Fibroblast-like synoviocytes, mitochondrial calcium uniporter, calcium

## Abstract

Rheumatoid arthritis (RA) is characterized by the aggressive migration and invasion of fibroblast-like synoviocytes (FLS) into cartilage and bone, a process that is significantly influenced by mitochondrial calcium uptake. This study highlights the critical role of mitochondrial calcium uniporter (MCU) in regulating FLS migration and mitochondrial dynamics, and its potential as a therapeutic target in RA. Notably, RA-FLS exhibited increased MCU expression and mitochondrial dysfunction compared to controls. Treatment with Ru360, a potent MCU inhibitor significantly reduced RA-FLS migration, calcium influx, and mitochondrial reactive oxygen species (ROS) levels, while restoring mitochondrial morphology and enhancing ATP production. Further analysis of MCU complex expression revealed elevated levels of MCU and other regulatory subunits (EMRE, MICU1, MICU2) in RA-FLS compared to controls, indicating mitochondrial dysfunction in RA. Mechanistically, MCU inhibition altered gene expression related to cytoskeletal dynamics, focal adhesion, and metabolic pathways. In human RA-FLS, MCU inhibition suppressed migration and invasion both in vitro and in vivo. The interaction between MCU and Miro1, essential for mitochondrial transport, was validated, and its disruption impaired FLS migration. Our findings highlight MCU as a promising therapeutic target to inhibit FLS migration and ameliorate RA progression.

## Introduction

Rheumatoid arthritis (RA) is a chronic, systemic autoimmune inflammatory joint disease. The global prevalence of RA is nearly 1% and It is more frequently observed in females than males. While biological DMARDs (bDMARDs) and targeted synthetic DMARDs (tsDMARDs) like Janus kinase (JAK) inhibitors, along with methotrexate (MTX), are currently used to reduce the inflammation, but still complete remission is yet to be achieved. Despite advancements, there is limited understanding of the disease pathogenesis particularly in the area of mechanism or genes responsible for the migration and invasion of fibroblast like synoviocytes (FLS). Aiming the invasion mechanism could generate new and better therapeutic outcomes.

In RA, the synovial joints are the primary target where the disease becomes established. A synovial joint is made up of two bone surfaces enclosed within a fibrous capsule lined with synovial membrane. Normally, the synovial membrane is 1-2 thin layer membrane composed of FLS. However, in RA, this membrane undergoes a dramatic transformation and become multilayered thick. This thickening contributes to joint stiffness and significantly reduces the quality of life **(Bottini and Firestein, 2013; Haroon et al., 2007).** FLS are the prime cells which surrounds the joint cavity and have been revealed to exhibit joint specific attributes imprinted during the initial phases of the joint development. Their extensive growth leads to the formation of an abnormal synovial tissue known as pannus. They start producing local inflammatory cytokines and enzymes, like matrix metalloproteinases (MMPs), which breakdown the extracellular matrix (ECM). This process encourages cells to move and invade, eventually causing the adjacent cartilage and bones to break down. **(Bustamante et al., 2017).** MMPs plays a crucial role in the establishment of RA. Besides cleaving extracellular components, they have the potential to directly activate signaling molecules such as tumour necrosis factor (TNF), which further contributes in the pathogenesis of the disease. FLS cells have the potential to express almost all MMPs except few **(Bian et al., 2023).** During the initial phases of the disease’s progression, several stimuli such as danger-associated molecular patterns (DAMPs), synovial citrullination, complement activation and antibodies activate the FLS cells. This activation leads to the release of inflammatory mediators and the recruitment of immune cells, which in turn creates an inflammatory microenvironment in the synovial space. Subsequently, this environment is characterized by high levels of pro-inflammatory cytokines, growth factors, and an infiltration of inflammatory immune cells, which further robustly activates FLS cells **(Muller-Ladner et al., 2000)**. Under these conditions, several signaling pathways become activated, which significantly impact cells’ metabolism and promotes their division, growth, and survival. Once activated, FLS cells undergo changes in their metabolism, gaining new metabolic profiles that are necessary for their proper function **(Guma et al., 2016).** In RA, FLS cells transformed into an aggressive and invading phenotype resembling that of tumor cells.**(Mousavi et al., 2021)**. Interestingly, *in vivo* migration of RA-FLS have been documented which clearly shows the metastasis of the disease from site of infection to the other anatomical locations **(Dang et al., 2019).**

Although, the biology of FLS is not completely understood but certain genes have been reported which reduces its invasiveness in *in vitro* conditions such as Huntingtin-interacting protein 1 (HIP1) **(Laragione et al., 2018),** transient receptor potential vanilloid subfamily, type 2 channel (TRPV2)**(Laragione et al., 2019).** Additionally, in rats, Cia5d has been found to be novel genetic regulator of the invasive properties of synovial fibroblasts **(Laragione et al., 2008).**

The literature strongly suggests the importance of increased intracellular Ca^2+^ concentration in the migration of various cell types including cancer cells. Mitochondria is popularly known to regulate the Ca2+ homeostasis via mitochondrial calcium uniporter (MCU) **(Tang et al., 2015).** The MCU protein complex localizes in the inner mitochondrial membrane and alone is essential and adequate for mitochondrial calcium uptake. A number of proteins contributes to the formation and regulation of this complex including MCU, EMRE, MCUR1, MICU1, MICU2 **(De Stefani et al., 2015)**. Studies show the correlation of mitochondrial calcium uptake via MCU, is responsible for the tumor growth and metastatic formation. Even more, silencing of MCU hampered cell motility and invasiveness of the cancer cells**(Tosatto et al., 2016).**

Apart from this, MCU has also been identified to regulate the mitochondrial dynamics. In order to maintain their shape, quality, distribution, size, and function, mitochondria undergoes orchestrated cycles of fission and fusion processes, a phenomenon termed as mitochondrial dynamics **(Chen et al., 2023)**. So, the balanced mitochondrial dynamics is crucial for the ideal function of mitochondria and cell fate **(Youle and van der Bliek, 2012)**. Evolving studies indicate that mitochondrial dynamics are important contributors to diverse cellular functions such as cell metabolism, migration, cell differentiation, apoptosis, and so on. Of note, the imbalanced mitochondrial dynamics is linked to a range of diseases, often characterized by impaired mitochondrial function and abnormal cellular fate **(Chen et al., 2023)**. Moreover, a study has demonstrated that Drp1-dependent mitochondrial fission is essential for the invasiveness of migrating cells because it redistributes mitochondria to the lamellipodia at the leading edge of breast cancer cells.**(Zhao et al., 2013).** More mitochondrial fission has been observed in MCU inhibited cells because Drp1, a crucial regulator of mitochondrial fission localizes on the outer mitochondrial membrane is partially regulated by ca2+ dependent signaling. In MCU deficient cells, cytoplasmic Ca2+ transients will be disturbed leading to altered Drp1 phosphorylation. Furthermore, MCU independently is unable to alter the cell proliferation, it requires Drp1 to influence the cell process **(Koval et al., 2019).** Interestingly, Drp-1 has been found to regulate cell migration via making mitochondria more fragmented in migrating cell. Furthermore, silencing of mitochondrial fusion proteins like Mfn-1 results in more cell migration **(Zhao et al., 2013).**

In this study, we investigated the role of MCU in RA pathogenesis. We hypothesized that inhibition of MCU might be instrumental for FLS cells migration and invasion to the surrounding tissues and could be a potential therapeutic target for clinical investigations.

## Materials and methods

### Patients

Patient samples were collected from the Department of Clinical Immunology and Rheumatology, Sanjay Gandhi Post-graduate Institute of Medical Sciences, Lucknow, Uttar Pradesh. The study was carried out in accordance with the Declaration of Helsinki. Patients were briefed on the aims of the study and gave their consent in writing. A DAS28 score of less than 2.6 indicates that the disease is in remission. Scores between 2.6 and 3.2 suggest low disease activity, while those between 3.2 and 5.1 represent moderate disease activity. A DAS28 score higher than 5.1 signifies high disease activity. The research was approved by the ethics review boards of both the Central Drug Research Institute (CDRI) and Sanjay Gandhi Postgraduate Institute of Medical Sciences (SGPGI).

In all patients, disease activity severity was assessed as described below:

DAS28 = 0.56 × √(TJC28) + 0.28 × √(SJC28) + 0.70 × in (ESR) + 0.014 × GH.

**PGA:** patient global assessment, **SJ**: swollen joint, **TJ**: tendon joint, RF: Rheumatoid Factor, **ESR**: Erythrocyte Sedimentation Rate, **DAS**: Disease activity score, **GH**: general health.

### Interpretation

DAS28 < 2.6: Remission

2.6 ≤ DAS28 ≤ 3.2: Low disease activity

3.2 < DAS28 ≤ 5.1: Moderate disease activity.

DAS28 > 5.1: High disease activity

### Cell culture

Primary RAFLS were isolated from synovial fluids and tissues after removing fat, fibrous membranes, and cartilage fragments. The synovial tissue was chopped into small pieces and digested overnight at 37°C with 1mg/mL type IV collagenase (Merck Millipore, USA) mixed in Dulbecco’s Modified Eagle Medium (DMEM). The resulting mixture was then filtered through a 70μm mesh cell filter (Merck Millipore, USA) into a 50mL tube and centrifuged at 250g for 10 minutes. The supernatant was discarded, and the cell pellet was rinsed with DMEM three times. The cell pellet was resuspended in DMEM containing 10% fetal bovine serum (FBS) and 100U/mL penicillin-streptomycin (Gibco, USA). The cells were cultured in 10cm culture dishes at a density of 2.5×10^4 cells/mm² in a humidified incubator (Thermo Fisher Scientific, USA) with 95% air and 5% CO2 at 37°C and were subcultured upon reaching confluence. Cells from passages 3 to 8 were used in the experiments.

### Animals

Male DBA/1J mice were acquired from the animal facility at the CSIR-CDRI in Lucknow, India. They were housed in a pathogen-free conditions with a 12-hour light/12-hour dark cycle. The animals were provided with chow and water ad libitum. Mice aged 6 to 8 weeks and weighing between 18 and 25 grams were used for the experiment. Mice were categorized into two groups. To understand the role of MCU in RA, CIA has been induced in the mice. To induce collagen-induced arthritis, bovine type II collagen (2 mg/ml) was mixed with an equal amount of CFA (2 mg/ml) to create an emulsion. Subsequently, on day 0, 150 µl of the emulsion will be injected intradermally at the base of the tail of DBA/1J mice. Twenty-one days after the initial primary immunization, mice will receive a boost with 100 µl of collagen emulsion prepared in IFA. After 40 days, mice were sacrificed, and FLS cells were collected from both control and RA mice. RA was assessed by monitoring the inflammation in the paws. The CDRI Institutional Animal Ethics Committee review board approved the animal studies.

### Xenograft model of rheumatoid arthritis

RA-FLS (5×10^5^) were transfected with *siScr* and *siMCU* for 24 hours. The cells were then injected intradermally into the backs of athymic nude mice (CSIR-CDRI, Animal facility). One day prior to FLS implantation, skin inflammation was induced by injecting complete Freund’s adjuvant (CFA) subcutaneously at a fixed distance (1.5 cm) from the site where RA-FLS would be injected. Five days later, skin tissue samples were collected from the RA-FLS implanted site and CFA-administrated site.

### siRNA transfection

RA-FLS were seeded in 12-well plate and reached 80% confluency by the day of transfection. Cells were treated with 100nM *siMCU*#1 (Invitrogen, USA) *siMCU*#2(santa cruz), or scramble *siRNA* (santa cruz) by Lipofectamine RNAiMAX transfection reagent (Invitrogen, USA) according to the manufacturer’s instruction. After the siRNA treatment, the cells were incubated at 37⁰C with 5% CO_2_ for 24 to 48 hours before being processed for further analysis.

### Immunohistochemistry

For immuno-histochemical staining, mice tissues were fixed in 10% neutral paraformaldehyde solution, followed by washing, dehydration and embedded in paraffin wax. 5-µm thick sections were taken on the slides. Sections were dewaxed in xylene and rehydrated followed by antigen retrieval using 10mMsodium citrate (pH:6) buffer. The tissue samples were incubated with bovine serum albumin (BSA) for an hour at room temperature to prevent unwanted binding. Afterwards, tissue sections were washed three times in PBST followed by incubation with HLA class I ABC antibody (1:10000, Proteintech) overnight at 4°C. The slides were rinsed three times with a phosphate buffer solution (PBST) and incubated with HRP-conjugated anti-mouse IgG (Peprotech) in a humid chamber for 2 hours. HLA class I-positive cells were detected by 3’3-diaminobenzidine tetrahydrochloride (DAB; Sigma). The slides were counterstained with hematoxylin. Images were captured on a Nikon Eclipse E200., Japan, light microscope at 200x magnification. The number of HLA-class I (+) cells, indicating RA-FLS, were analyzed using ImageJ Fiji software.

### Western Blotting

Tissue and cell proteins were extracted with RIPA lysis buffer and a 1% protease inhibitor cocktail (HiMedia). The lysate was centrifuged at 12,000 rpm for 30 min followed by a collection of supernatant and the amount of protein in it was measured using a Bradford assay. Next, an appropriate volume of 5x loading buffer was added, samples were boiled at 95⁰C for 10 min. Protein samples were subjected to 10% SDS-polyacrylamide gel electrophoresis, followed by transfer to PVDF membranes. After the membranes were blocked using 5% Bovine Serum Albumin (BSA) in TBS-Tween, they were then left to incubate at 4°C overnight with primary antibody targeting MCU (Proteintech, 1:3000,Catalog no: 26312-1-AP), β-tubulin (Abclonal, 1:10,000, Catalog No.: AC021). Following a wash with TBS-Tween20, the membranes were treated with secondary goat anti-mouse (Affinity, USA, 1:5000) or anti-rabbit (Affinity, USA, 1:5000) antibodies. The protein bands were then visualized using ECL chemiluminescent detection solutions (Bio-Rad).

### Mitochondrial ultrastructure with transmission electron microscopy

RAFLS cells were seeded up to 90% confluency in the presence and absence of Ru360. Alterations in mitochondrial ultrastructure due to Ru360 was then analyzed using TEM. To check the effect of Ru360, human RAFLS were fixed with 2.5% glutaraldehyde in phosphate buffer for 24 h, followed by rinsed with phosphate buffer, afterwards fixing fixed in osmium tetroxide, and encapsulated in agarose. The sample was cells were dehydrated using a series of ethanol solutions with increasing alcohol concentrations. Afterward, it was encased, cells were embedded in epoxy resin and cured at a temperature of 60⁰C for 24 hours. 50–70 nm ultrathin sections were taken obtained using a LEICA EM UC7 ultramicrotome, which were then collected on copper grids and stained with uranyl acetate and lead citrate. The grids were then studied, observed at an accelerating voltage of 100 kV accelerating voltage using under a JEOL JEM-1400 TEM equipped with a Gatan bottom-mounted Orius CCD camera.

### ROS measurements

Mitochondrial ROS were measured using MitoSOX, a fluorescent dye (Life Technologies). Cells were exposed to Ru360 (at a concentration of 3 uM) for 24 hours, with or without the addition of TNF (at a concentration of 20 ng/ml) for the 24 hours. Following this treatment, cells were incubated with 100 nM MitoSOX for 30 minutes at 37⁰C, washed, and then analyzed for fluorescence intensity using a flow cytometry (FACSAria II cell sorter, BD Biosciences).

### Wound healing assay

The cells were allowed to grow to 80–90% confluency in 12-well plates. A 200 μl sterile pipette tip was used to create a central linear wound, followed by 2 to 3 times washing of wells with PBS to remove any floating cells. The wound closure rate was assessed by measuring the distance from the wound edge to the original wound site after 24 hours. Each experiment was performed from the three patients’ RA-FLS cells.

### Matrigel invasion assay

The Corning BioCoat Matrigel Invasion Chamber assay system (USA) was used to study RAFLS invasion, according to the manufacturer’s instructions. After transfection with scrambled *siRNA* and *siMCU*#1 (ThermoFisher Scientific, Silencer™ Pre-Designed siRNA, Catalog number: AM16708), *siMCU*#2 (SANTA CRUZ BIOTECHNOLOGY, INC., CCDC109A (E-9): sc-515930) for 24hrs, empty vector and pcDNA3.1/C-(K)-DYK MCU plasmid for 24 hours. RA-FLS were permitted to invade the matrigel invasion chamber for a further 24 h in the presence of incomplete DMEM and TNF (20ng/ml) in the lower surface. Similarly, RAFLS cells were treated with or without Ru360 for 24h followed by migration in matrigel invasion chamber in the presence of incomplete DMEM and TNF (20ng/ml) in the lower surface. The non-invading cells were gently wiped away using a cotton swab. The remaining cells, attached to the underside of the membrane, were then stained with crystal violet. For quantification, cells were counted in three randomly selected fields.

### Human Rac1 Analysis

Rac1 level was assayed using Human Rac1 (Ras related C3 Botulinum Toxin Substrate) ELISA kit from MyBioSource.com as per manufacturer instructions. Human RA-FLS were stimulated with TNF (20ng/ml) post transfection of siRNA (scrambled and siMCU#2), followed by lysis in RIPA lysis buffer and supernatants were subjected for Rac1 assay.

### Matrix metalloproteinase activity analysis

MMP activity level was assessed using MMP assay kit from ANASPEC in human RA-FLS cell lysates as mentioned in the product datasheet. RA-FLS were stimulated with TNF (20ng/ml) post transfection of siRNA (scrambled or siMCU#2) followed by lysis in RIPA lysis buffer and the supernatant was added in 1:1 ratio with MMP colorimetric substrate. After 12hrs the end point absorbance is taken at 412nm.

### RNA extraction, reverse transcription, and quantitative real-time PCR

Total RNA was extracted from RA-FLS cultures utilizing trizol reagent (Thermo Fisher Scientific, USA). Subsequently, the RNA underwent reverse transcription into cDNA using the ReverTra Ace qPCR RT Master Mix with gDNA remover kit as per the protocol provided by the manufacturer (Toyobo, Japan). The q-PCR analysis was carried out with the SYBR Green Realtime PCR Master Mix kit (Toyobo, Japan) on a Quant Studios 7 Flex real-time PCR system (Applied Biosystems, USA). The sequences of the primers are provided in Table 1.

### Statistics

GraphPad Prism (https://www.graphpad.com/features) was used to perform statistical analysis. p value was determined by 2-tailed student’s test or 1-way/2-way ANOVA. Results were considered statistically significant if the p-value was less than 0.005. Data is presented as the average value (mean) with standard error of the mean (SEM). The number of samples and independent repeats are provided in the figure legends. ImageJ and fiji was used for the quantification of microscopy.

### Metabolite Extraction

All the cell lines samples were processed in triplicate. In order to extract metabolites, two million cells were collected. At first, 500 μL of cold MeOH (–20°C) was used to resuspend the cell pellets. Then, 30 μl of ribitol solution (0.2 mg/ml in water) was added to every sample as an internal standard. The samples underwent rapid freezing in liquid nitrogen and then vortexed for 2 minutes. After thawing at 37°C, the samples were centrifuged at 800g for 5 minutes at 4°C. The supernatant was thereafter transferred to a fresh centrifuge tube. Using 500 μL of a chilled MeOH solvent, the freeze-thaw cycle was repeated once again. Lastly, the sample was centrifuged at 15,000 g for 2 minutes after being mixed with 250 μL of ice-cold water for the freeze-thaw extraction procedure. The collected supernatant of all the samples were dried by vacuum centrifugal evaporation. Thereafter, 20 μL of MOX reagent, containing 20 mg/mL of methoxyamine-hydrochloride in dry pyridine, was added to the dried samples. A shaking dry bath (Thermomixer, Eppendorf) that was digitally heated to 30°C and 1100 rpm was then used for an incubation period of 90 minutes. The derivatization process began with the addition of 80 μL of N-Methyl-N-(trimethylsilyl) trifluoroacetamide with 1% trimethylchlorosilane (MSTFA + 1% TMCS), and was continued for 30 minutes in a Thermomixer set at 60°C.

### GC-MS analysis

The prepared samples were analyzed using a gas chromatography-mass spectrometry (GC-MS) system. To ensure reliable results, a standard mixture was analyzed between each sample to check instrument performance and data accuracy. 1 µl of the samples was introduced in splitless mode through a Triplus 100 autosampler (Thermo Scientific) into a Trace 1300 gas chromatograph, which was equipped with a TSQ 8000 mass spectrometer. The metabolites present in the samples were separated using a TraceGOLD TG-5MS column (Thermo Scientific), which features a diameter of 0.25 mm, a film thickness of 0.25 µm, and a total length of 30 m. Samples were transported and analyzed using high-purity helium and argon gases, maintaining a flow rate of one milliliter per minute. The point where the sample is injected was heated to 200⁰C, while the connecting tube and the part that ionizes the sample were both set to 250⁰C. The oven initially started at a temperature of 50⁰C and was kept constant for one minute. Subsequently, the temperature was steadily increased to 100⁰C at a rate of 6⁰/minute. This was followed by another increase to 200⁰C at a slower rate of 4⁰ per minute. Finally, the temperature was rapidly raised to 280⁰C and maintained at that level for three minutes. All samples were analyzed in full scan mode, covering range from m/z 60 to 650. The resulting data was collected for further examination.

### Data Processing

The raw data was processed using MS-DIAL software version 4.90. This involved identifying significant peaks, separating overlapping signals, aligning spectral data, and assigning potential identities to the detected compounds. The electron ionization mass spectrometry (EI-MS) spectra were analyzed and the metabolites were categorized using Level 2 (authentication via spectral databases) and Level 3 (tentative identification based on spectral similarity to known compounds within a chemical class), in accordance with the metabolomics standards initiative (MSI) guidelines for metabolite identification. The identified metabolites were subjected to statistical analysis using MetaboAnalyst 5.0 software. To prepare the data for analysis, the values were adjusted (normalized), transformed using a cube root function, and scaled using a Pareto method.

## Results

### Comparative analysis of RA-FLS migration in RA and control DBA1/J mice

FLS cells are the principal cellular component of invasive pannus in RA, with the potential to migrate and invade nearby cartilage and bone by secreting proteases. To study the migratory properties, we isolated FLS cells from the synovial joint of control and collagen-induced arthritis (CIA) RA mice (**Figure 1A**). Recent studies have highlighted the importance of mitochondrial calcium in cell migration (**Paupe and Prudent, 2018**). To explore this, we performed a wound-healing assay and attempted to inhibit their migration using Ru360,a, a potent inhibitor of calcium uptake via MCU.

**Figure 1:**
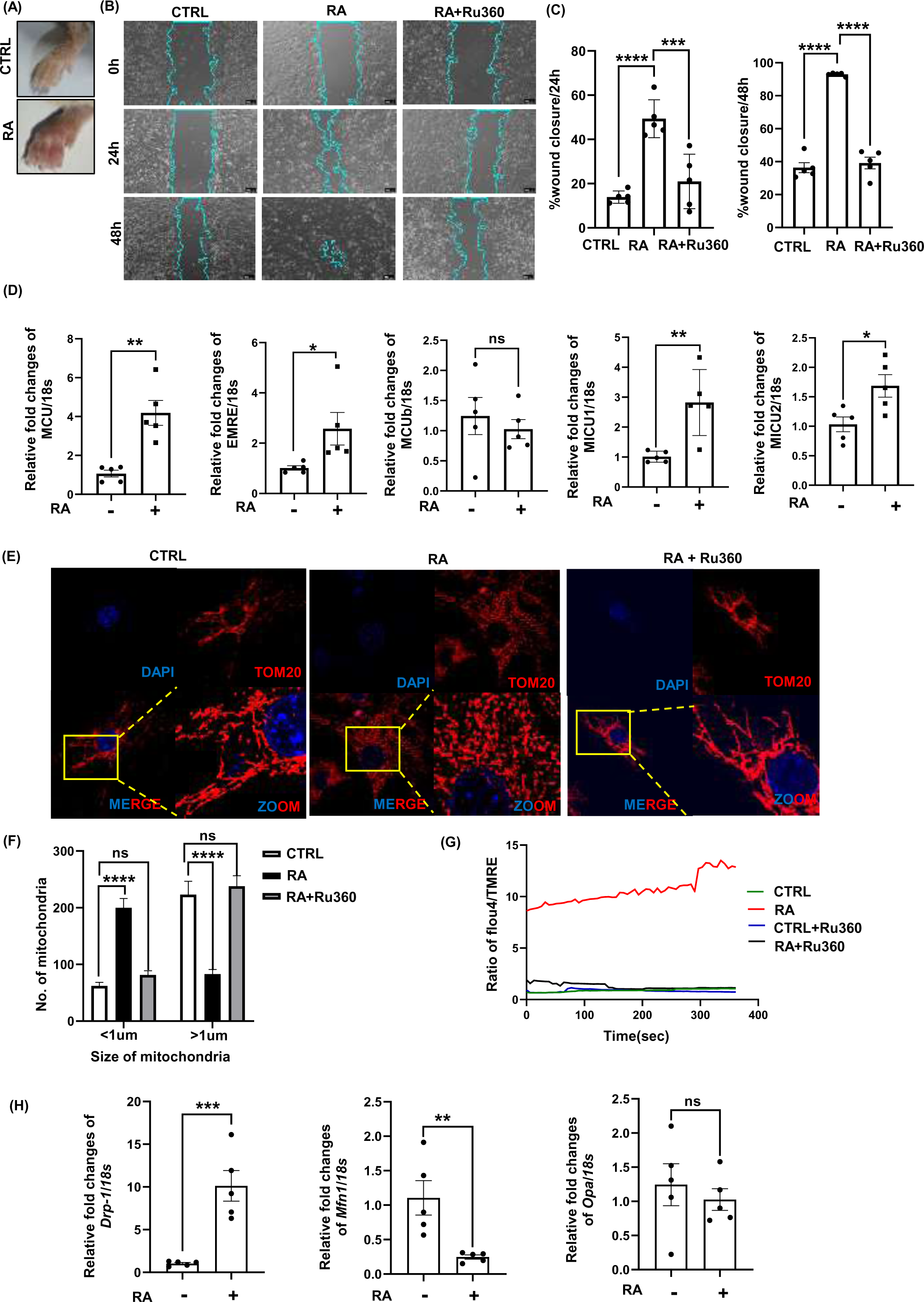
Role of MCU in RAFLS cells. (**A**) Comparative Analysis of paw Inflammation in control and RA mice. Control mice show no sign of inflammation while RA mice have significant swelling and redness in the paw. **(B)** Representative images of wound healing in control (CTRL) and rheumatoid arthritis (RA) RAFLS cells over time, along with RA cells treated with Ru360 (a mitochondrial calcium uniporter inhibitor), at 0 hours (initial wound) and 24-and 48-hours post-wounding. The wound edges are outlined in blue to highlight the extent of closure over time. Images were taken at 4x magnification. **(C)** Quantitative analysis of wound closure percentage in CTRL, RA, and treated RA cells treated with Ru360. The treatment groups show varying degrees of wound healing improvement compared to untreated RA cells. **(D)** Expression of MCU complex genes. The bar graphs display the relative expression of various mitochondrial calcium uniporter (MCU) complex components in control and CIA-DBA1J mice FLS cells. **(E)** Effect of Ru360 on mitochondrial morphology in RA fibroblasts. Representative confocal microscopy images of FLS cells stained with DAPI (blue) for nuclei and TOM20 (red) for mitochondria. Mitochondria appear highly fragmented in RAFLS compared to control FLS cells. Cells treated with Ru360 (a mitochondrial calcium uniporter inhibitor) display increased mitochondrial elongation similar to control FLS cells. **(F)** Quantification of mitochondrial size. The bar graphs display the relative size of mitochondria (<1 µm, and >1 µm,) between control (CTRL), rheumatoid arthritis (RA), and RA fibroblasts treated with Ru360 (RA+Ru360). The number of fragmented mitochondria (size <1um) was higher while elongated mitochondria (size >1um) was significantly lower in RAFLS compared to control FLS. Treatment with the MCU inhibitor Ru360 normalized the mitochondrial size in RAFLS cells to levels comparable to the control group, showing no significant difference (ns). **(G)** Dynamics of calcium influx in control (CTRL), rheumatoid arthritis (RA), control FLS treated with Ru360, RA treated with Ru360 (RA + Ru360). The ratio of Fluo-4 (calcium indicator) to TMRE (mitochondrial membrane potential indicator) fluorescence was measured over time. RAFLS cells exhibited a significant and sustained increase in calcium influx compared to control cells. Treatment of RAFLS with the MCU inhibitor Ru360 significantly reduced calcium influx, normalizing it to levels comparable to the control group. **(H)** Relative fold changes of mRNA expression for mitochondrial fission (*DRP1*) and fusion (*MFN1*), (*OPA1*) normalized to 18S rRNA in control and mice RAFLS. Data represent the mean ± SEM of three independent experiments. Statistical significance was determined using ANOVA. *\*p < 0.05, **p < 0.01, ***p < 0.001, ****p < 0.0001, ns = non-significant*. Scale bars indicate 10 μm.

In the wound healing assay, we observed significant differences in wound closure rates among control FLS, RA-FLS and RA-FLS treated with Ru360. Interestingly, the FLS from the synovial joints did not exhibit migratory characteristics, whereas RA-FLS cells showed strong migratory properties, as depicted in (**figure 1B**). Moreover, RA-FLS treated with inhibitor demonstrated a notable reduction in cell migration compared to untreated RA-FLS (**figure 1C**). These results indicate that administration of Ru360 effectively blocks the calcium uptake via MCU, thereby reducing cell migration.

### MCU complex expression in RA and control FLS

As the inhibition of MCU alters cellular migration, we were prompted to further investigate the expression of MCU complex genes. We compared the expression levels of *MCU* complex genes in FLS isolated from control and RA mice (**figure 1D**). Interestingly, we observed an elevated expression of MCU in RA compared to control FLS cells. Additionally, the expression of other regulatory and calcium sensor genes, including EMRE, MICU1, and MICU2, were also increased. Of note, the negative regulatory subunit (MCUb) exhibited no significant changes. Taken together, these findings suggest that the expression of MCU complex genes is augmented in RA, indicating a potential role of mitochondrial dysfunction in the disease.

### MCU regulates microdynamics

Our initial findings revealed an upregulation of MCU complex genes in RA FLS cells, indicating potential mitochondrial dysfunction. Moreover, previous studies have documented the role of calcium in regulating mitochondrial morphology(**Kristal and Dubinsky, 1997**),(**Dubinsky and Levi, 1998**), leading us to further examine the mitodynamics using confocal microscopy. We observed that RA-FLS cells exhibited fragmented mitochondria compared to the elongated mitochondria in control cells. Interestingly, treatment with an MCU inhibitor restored mitochondrial morphology to a state resembling that of control cells as depicted in **figure 1E,1F**. These results suggest that MCU-mediated calcium dysregulation may contribute to the observed mitochondrial fragmentation in RA FLS cells, highlighting a potential link between MCU and mitochondrial dynamics in the pathogenesis of rheumatoid arthritis.

Next, to confirm the above results we performed the mitochondrial calcium influx assay using calcium indicator Fluo-4 and the mitochondrial membrane potential indicator TMRE. We observed a marked increase in the calcium influx in RAFLS compared to the control FLS cells, indicating dysregulation of calcium homeostasis in the RA-FLS cells. Post-treatment with the MCU inhibitor, Ru360, the calcium levels in the RA-FLS cells were restored to levels comparable to those observed in the control FLS cells as depicted in **figure 1G**.

Furthermore, previous studies have established that DRP-1 induces increased mitochondrial fission, thereby promoting inflammation in rheumatoid arthritis fibroblast-like synoviocytes (**Wang et al., 2020**). To corroborate our findings, we investigated mitochondrial dynamics by assessing fission and fusion markers using RT-PCR. As anticipated, we observed elevated expression of the fission marker, Drp-1, and decreased expression of the fusion marker, MFN-1 (**figure 1H**).

### MCU regulates migration and invasion of human RAFLS

These findings strongly support the hypothesis that MCU plays a critical role in the pathogenesis of RA. To understand more about the role of MCU in disease pathology, we aimed to study human FLS isolated from synovial joints of rheumatoid arthritis patients. We collected synovial cells from eight patients with rheumatoid arthritis, and for each patient, the DAS28 score was determined, as illustrated in **figure 2A**. Initially, we assessed the expression of MCU using western blot **analysis from RA-FLS isolated from five patients**. The results, shown in **figure 2B**, indicated the presence of MCU expression. The regulation of cell migration, a critical factor in metastasis, has been increasingly recognized to be influenced by mitochondrial calcium. In this context, we attempted to inhibit mitochondrial calcium uptake using Ruthenium red (RR) and Ru360 (Ruthenium 360) at a concentration of 3 μM. Similar to what we observed in mice, we found a reduction in the migration of human RA-FLS as shown in **figure 2C-2F.** Although both inhibitors were effective in reducing migration to some extent, but RR was found to be toxic to the cells (**figure 2G**), leading us to choose Ru360 instead. To further evaluate the invasive potential of human RA-FLS cells, an invasion assay was conducted in the presence and absence of the Ru360. The results from the invasion assay corroborated the findings from the wound healing assay, showing a marked reduction in invasion after treatment with Ru360 (**figure 2H,2I**).

**Figure 2:**
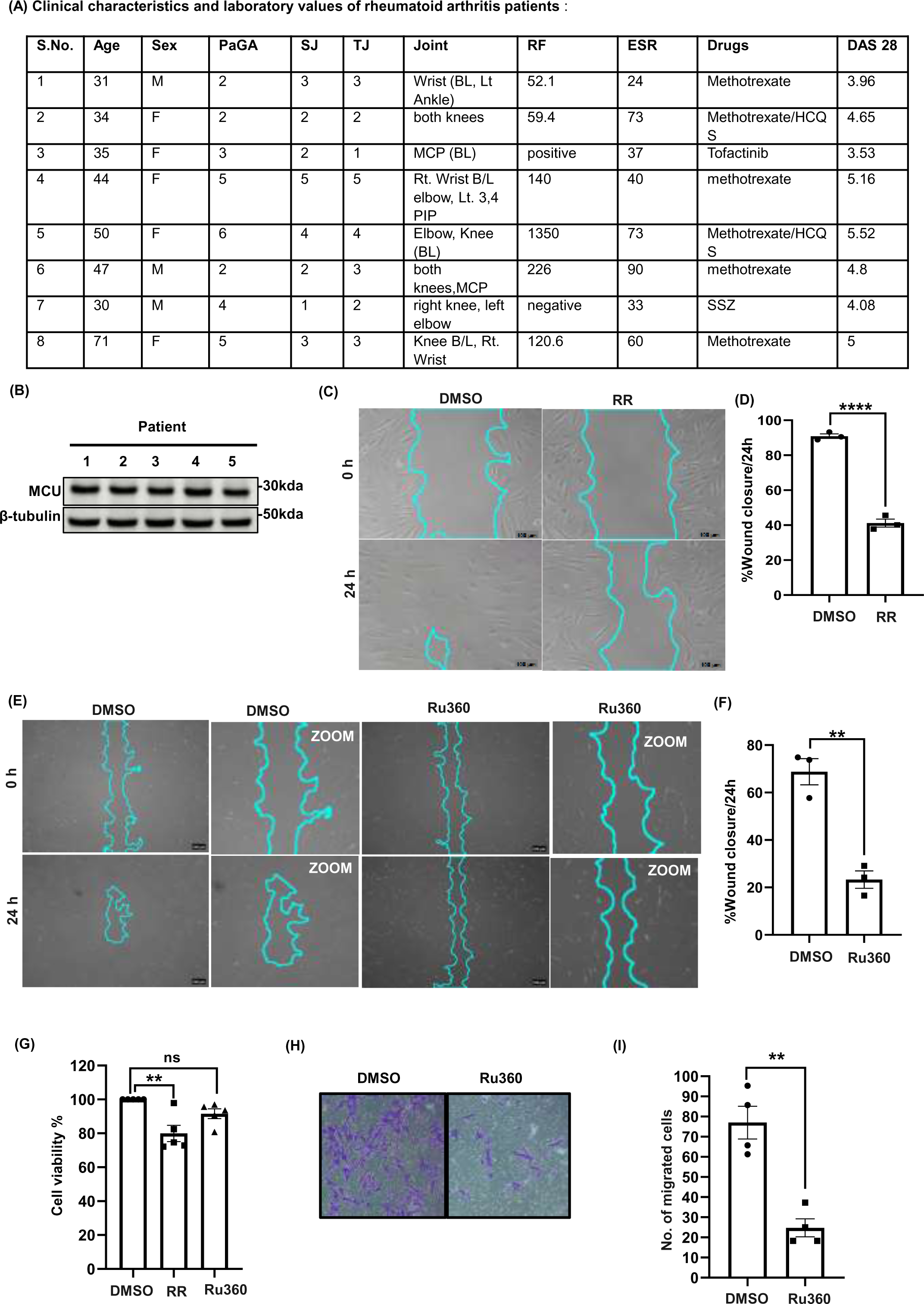
Impact of Ru360 on human RA-FLS. (**A**) The table represents the demographic and clinical data of rheumatoid arthritis (RA) patients. **(B)** Western blot analysis of MCU protein expression from five RA patient’s RA-FLS are shown. Beta-tubulin was used as a loading control. **(C-F)** A wound healing assay was performed from freshly isolated 80-90% confluent human RA-FLS (n=3) with and without Ruthenium Red (RR) and Ru360, mitochondrial calcium uniporter (MCU) inhibitors. Representative images depict wound closure at 0 and 24 hours. The bar graph quantifies the percentage of wound closure over 24 hours. **(G)** A cell viability, MTT assay was performed with Ruthenium Red and Ru360 treatments. The bar graph quantifies the percentage of cell viability compared to the DMSO control. **(H)** Representative images of cell invasion assay in the absence (–Ru360) or presence (+Ru360) of Ru360. **(I)** Quantification of number of cells invaded through matrigel treated with DMSO (control) or Ru360. The inhibitor showed the significant reduction in the invasion of human RA-FLS. Data show mean ± SEM. *(** p < 0.01; *** p < 0.001, **** p < 0.0001),* analyzed using t-test

To ensure the specificity of our findings, we knocked down MCU using two *siRNA,* (**figure 3E**) and repeated the wound healing and invasion assays. *SIRNA#1* had three isoforms namely, isoform 1, isoform2 and isoform 3, while siRNA#2 was highly specific Consistently, silencing MCU with *siRNA* led to a significant decrease in the migration (**figure 3A, 3B**) and invasion abilities (**figure 3C, 3D**) of RA-FLS cells, corroborating the results obtained with Ru360 inhibition. These findings highlight the importance of MCU in regulating the migration and invasion of FLS cells in RA. Next, we sought to further validate the role of MCU in cell invasion by overexpressing the MCU gene in human RA-FLS. Subsequently, we performed the same invasion assay to assess the impact of MCU overexpression on cell invasion. We observed a marked increase in the invasion of RA-FLS compared to the empty vector (**figure 3F,3G**), reinforcing the idea that MCU might have the potential to regulate cell invasion. Moreover, to ensure the accuracy and specificity of genetic constructs and siRNAs potential, we first confirmed the overexpression (O.E) and knockdown (KD) of MCU first in human-RA-FLS cells, as depicted in **figures 3H, 3I.**

**Figure 3:**
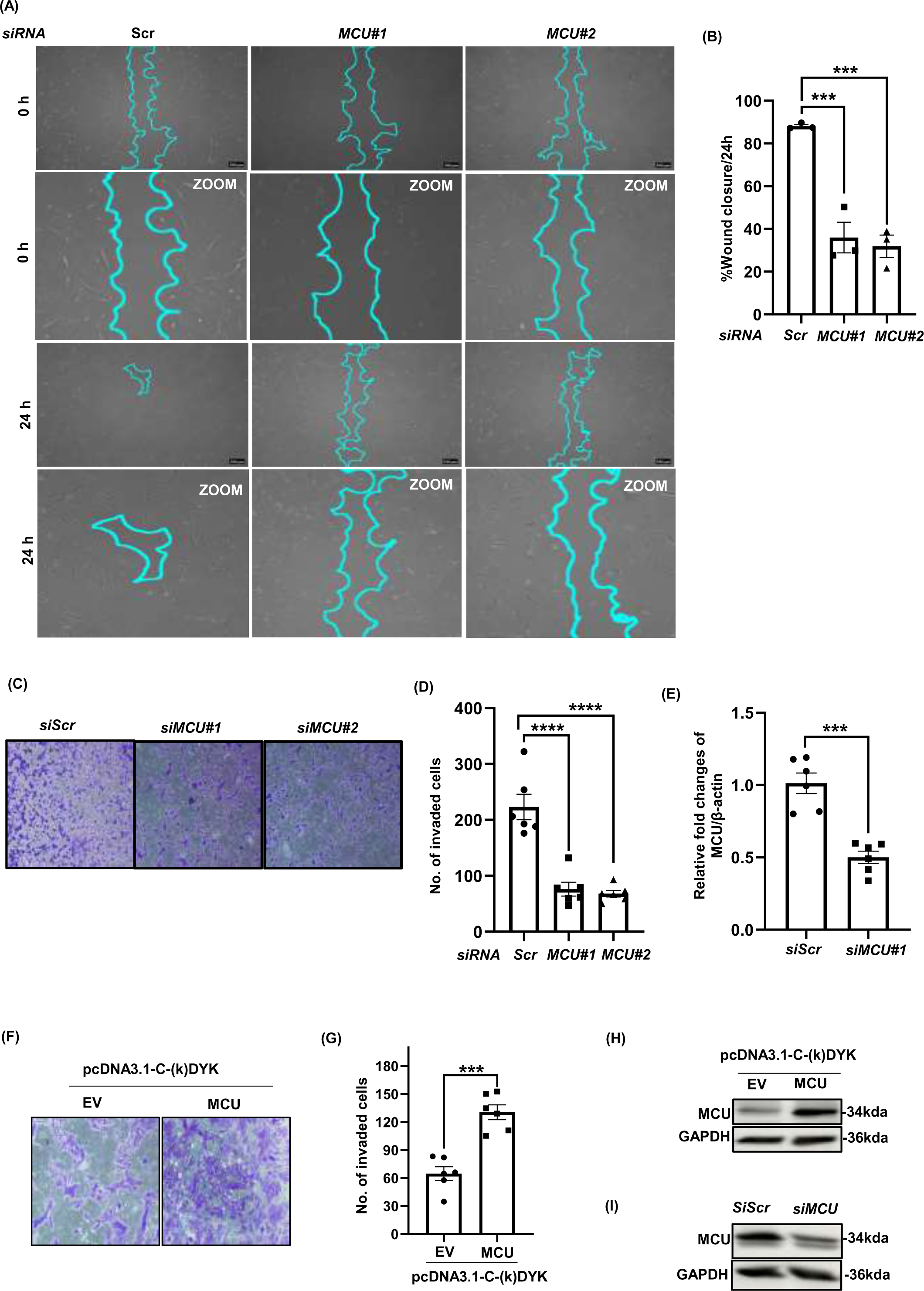
MCU Modulation affects migration and Invasion in RA-FLS cells:. (**A**) An additional wound healing assay was carried out with *siRNA* targeting MCU with 2 different *siMCU#1, siMCU#2* and compared to *scrambled siRNA* (*Scr*). Representative images illustrate wound closure at 0 and 24 hours. **(B)** The bar graph quantifies the percentage of wound closure over 24 hours. **(C)** Representative images of cell invasion through matrigel-coated membranes in cells transfected with *scrambled siRNA* (*siScr*) and *siRNA* targeting MCU (*siMCU#1, siMCU#2*) are shown. **(D)** Quantification of invaded cells invaded through matrigel are presented. **(E)** *MCU mRNA* expression was reduced by *MCU siRNA* transfection for 48h, as revealed by real-time PCR. Quantification of knockdown efficiency relative to β-actin, normalized to the control group. Data are represented as mean ± SEM (n=6 patients). Data show mean ± SEM. *(** p < 0.01; *** p < 0.001, **** p < 0.0001)*, analyzed using t-test. **(F)** Representative images of cell invasion through matrigel-coated membranes in cells transfected with empty vector (*EV*) or *pcDNA3.1-C-(k)DYK* (MCU overexpression). **(G)**The bar graph illustrates the enhanced invasion upon MCU overexpression. **(H-I)** Western blot analysis of MCU protein expression. MCU overexpression in cells transfected with *pcDNA3.1-C-(k)DYK* compared to empty vector (*EV*) control. MCU knockdown in cells transfected with *siRNA* targeting MCU (*siMCU*) compared to *scrambled siRNA* (*siScr*) control. GAPDH was used as a loading control.

### MCU inhibition downregulate the cytoskeleton dynamics

To gain deeper insights into the molecular mechanisms underlying the role of MCU in the RA-FLS migration, we performed an unbiased RNA sequencing (RNA-seq) of human RA-FLS +/− TNF and RA-FLS+TNF +/− Ru360 (**figure 4A)**. We wanted to understand how MCU influences the cell migration and invasion. We choose TNF because it effectively activates FLS cells and triggers them to produce pro-inflammatory cytokines, that closely resemble the inflammatory milieu of RA (**Koedderitzsch et al., 2021**). Principal component analysis (PCA), volcano plot and heatmap of the RNA sequencing data validated the consistency of the biological replicates for each condition as shown in **figure 4B**. We identified a set of 10228 genes showing differential expression between RA FLS cells with and without TNF, and 11,836 genes showing differential expression between RA FLS cells treated with TNF, with and without Ru360. Among these, 599 genes were upregulated and 357 were downregulated in response to TNF, whereas 381 genes were upregulated and 1,124 were downregulated in response to the combination of TNF and Ru360. In our transcriptome analysis, KEGG pathways highlighted the marked downregulation of genes involved in the actin cytoskeleton and focal adhesion pathways upon MCU inhibition. This downregulation has profound implications for cell motility, adhesion, and invasive potential. Key proteins involved in orchestrating these processes include Rac, PAK, F-actin, LIMK, Raf, MEK, Asef, Arp2/3 and so on.

**Figure 4:**
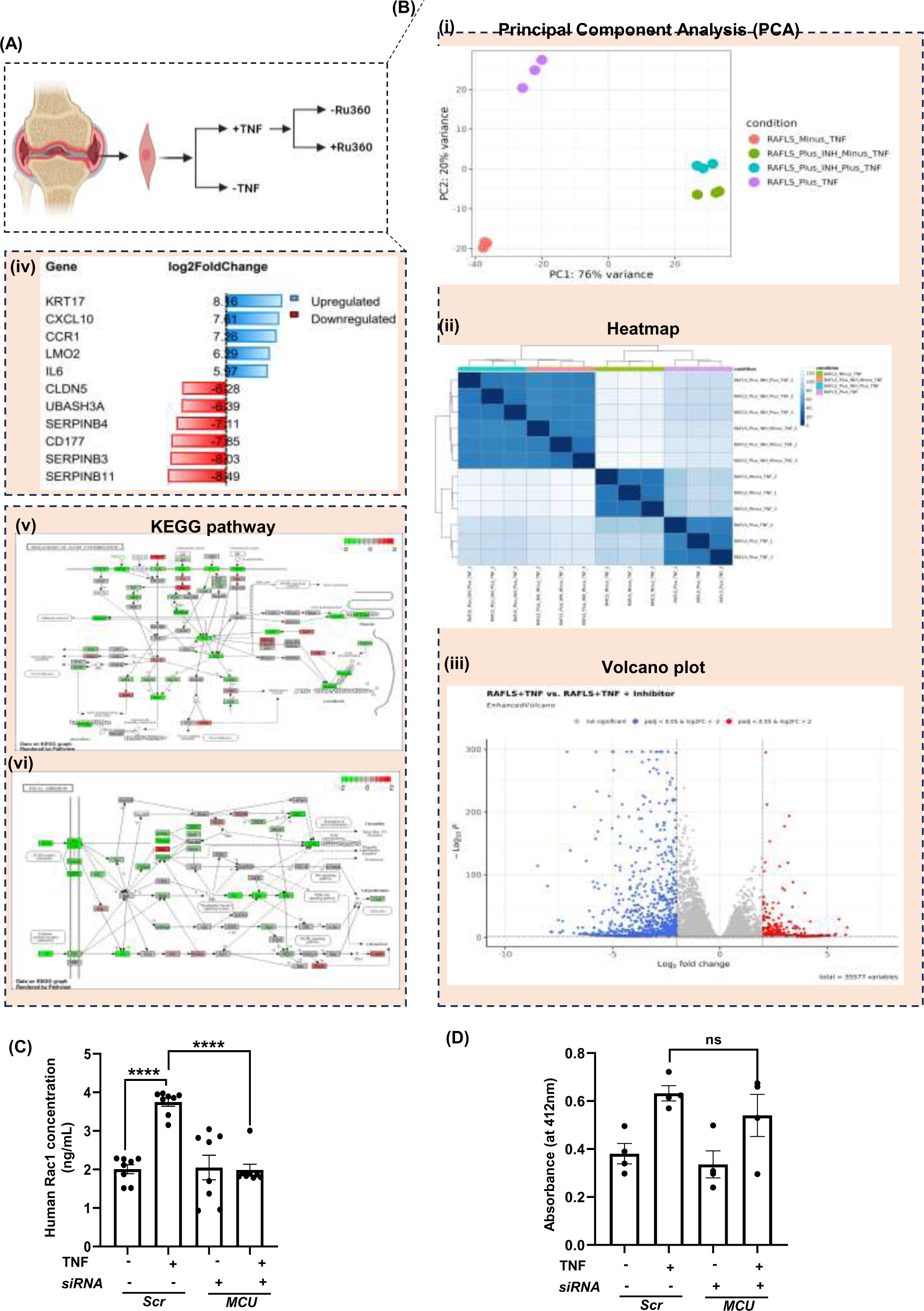
Comprehensive analysis of gene expression and functional pathways in RA-FLS. (**A**) Schematic representation of experimental conditions. RAFLS cells were treated with TNF, Ru360, or a combination of both. **(B) (i)** Principal component analysis (PCA) of gene expression data. Each dot represents a sample. **(ii)** Heatmap of differentially expressed genes (log2 fold change) across different conditions. Gene clusters are shown in rows, and conditions are shown in columns. **(iii)** Volcano plot showing differentially expressed genes between RAFLS+TNF and RAFLS+TNF+Inhibitor conditions. Red dots represent significantly upregulated genes, blue dots represent significantly downregulated genes. **(iv)** Log2Fold Change of differentially expressed genes. The figure displays the log2 fold change of differentially expressed genes. Positive values indicate upregulated genes (blue bars), while negative values indicate downregulated genes (red bars). **(v)** This figure depicts the KEGG pathway map for the regulation of the actin cytoskeleton. The pathway highlights key proteins and genes involved in actin polymerization, depolymerization, and regulation**. (vi)** The KEGG pathway map for focal adhesion. The pathway highlights key proteins and genes involved in cell-matrix interactions and signaling. Green and red boxes indicate upregulated and downregulated genes, respectively. **(C)** Rac1 release is low in *siMCU* treated cells as compared to scrambled cell lysate. Human RA-FLS were transfected with *siMCU*, stimulated with TNF and analyzed for RAC 1 by ELISA (n=8). Mean ± SEM; Statistically significant differences between indicated groups are marked with asterisks*(****p<0.0001)*; as determined by one-way ANOVA. **(D)** MMP activity absorbance appears to be low in *siMCU* treated cells as compared to scrambled cell lysate and the absorbance in cells without *siMCU* is significantly high. Human RA-FLS were transfected with *siMCU*, stimulated with TNF and analyzed for MMP activity (n=4). Mean ± SEM; Statistically significant differences between indicated groups are marked with asterisks *(* p<0.05)*; as determined by one-way ANOVA.

After inhibiting MCU, we observed the downregulation of these key genes, which might be responsible for the reduction in cell migration. One of the most impacted pathways involves Rac1, a crucial regulator of the actin cytoskeleton. Rac1 orchestrates the formation of lamellipodia, which are essential for cell movement. The downregulation of genes associated with Rac1 signaling indicates a diminished ability of cells to reorganize their actin cytoskeleton in response to migratory cues. This disruption hampers the polymerization of actin into filamentous actin (F-actin), a process critical for the formation of protrusive structures like lamellipodia and filopodia **(Ma et al., 2023)**. Similarly, PAK, a downstream effector of Rac1, was also affected. PAK plays a role in stabilization of actin filaments by phosphorylating LIM kinase (LIMK), which inactivates the actin-severing protein cofilin. The reduction in PAK and LIMK signaling pathways probably leads to a decrease in the stability of actin filaments, which in turn affects the cell’s capacity to create protrusions essential for its movement **(Zeng et al., 2020)**. Additionally, the pathway known as Raf-MEK-ERK, which is involved in the regulation of gene expression concerning cell mobility, was inhibited. This downregulation may contribute to the reduced expression of genes involved in cytoskeletal remodeling and focal adhesion turnover, further compromising cellular motility. Asef, a guanine nucleotide exchange factor (GEF) that activates Rac1, also showed reduced activity. Asef’s downregulation likely leads to impaired Rac1 activation, further limiting the formation of lamellipodia and reducing cell migration and invasion **(Itoh et al., 2008)**. Finally, the Arp2/3 complex, which plays a vital role in actin nucleation and branching, was downregulated **(Papalazarou and Machesky, 2021)**. This likely compromises the cell’s ability to form the branched actin networks essential for lamellipodia formation and effective cell migration. These combined disruptions in Rac1 signaling, actin stabilization, and cytoskeletal organization highlight the critical role of MCU in maintaining the cellular machinery required for migration and invasion.

### Increased mitochondrial calcium induces Rac1 activation

However, it remains unclear whether mitochondrial dysfunction is the primary cause of the disease or a secondary effect. The connection between mitochondrial dysfunction and cell migration is still lacking. Currently, there is a growing focus on calcium-dependent RAC1 activation in ongoing research **(Price et al., 2003**). Of note, Rac regulates the cytoskeletal organization of the lamellipodia formation which is coordinated with the process of cell migration **(Hall and Nobes, 2000)**. To decipher the role of mitochondrial calcium-dependent Rac function in migrating cells and to support the observations from RNA-seq, we have conducted Rac1 assay in the cell lysate of the human RA-FLS cells. We knocked down MCU using siRNA, followed by stimulation with TNF, and then measured RAC1 activity. We observed a significant reduction in RAC1 activity (**figure 4C**) after MCU inhibition compared to the control without MCU inhibition.

In our transcriptome analysis, we observed the downregulation of MMPs. Next to know whether MCU inhibition would impact the release of MMPs—proteins crucial for breaking down the extracellular matrix, a crucial first step in cell migration which is essential for the degradation of ECM the initial step to migrate. We again silenced the *MCU* via siRNA, followed by TNF stimulation. However, when conducting the MMP assay, we measured overall MMP levels rather than specific ones, we found no significant change in total MMP activity. (**figure 4D**). So, taken together, our findings suggest that MCU inhibition, significantly reduces Rac1 activity, which may influence cytoskeletal organization and cell migration. Although our transcriptome analysis indicated the downregulation of specific MMPs, the lack of significant change in total MMP activity following MCU inhibition suggests that the impact may be on specific MMPs rather than the overall MMP pool.

### MCU attenuation significantly reduces mitoROS

It is well established that mitochondrial ROS production plays a role in cellular migration(**Tosatto et al., 2016)**. Interestingly, various antioxidant molecules including MARVEL(MitoQ), Curcumin etc**.(Promila et al., 2023)** has been demonstrated to inhibit cell migration both *in vitro* and *in vivo* **(Porporato et al., 2014).** Consistent with these findings, we proposed that inhibiting calcium uptake could lead to a reduction in mitoROS levels. Accordingly, we pretreated human RA-FLS cells with Ru360 under TNF-stimulated conditions. We observed a significant reduction in the mitochondrial ROS production (**figure 5A**), suggesting, Ru360 can be the potent inhibitor to reduce the mitochondrial ROS in the inflammatory condition. These results prompted us to investigate whether the inhibitor has any effect on mitochondria health. We examined mitochondrial damage using MitoTracker Green and Red in human RA-FLS cells after treatment with Ru360 under TNF-stimulated conditions and compared the results with untreated cells. Interestingly, this mitochondrial damage was lessened when TNF-stimulated cells were treated with Ru360 (**Figure 5B**). Taken together, our findings suggest that blocking MCU likely prevents an excessive buildup of calcium, thereby protecting mitochondria from dysfunction

**Figure 5:**
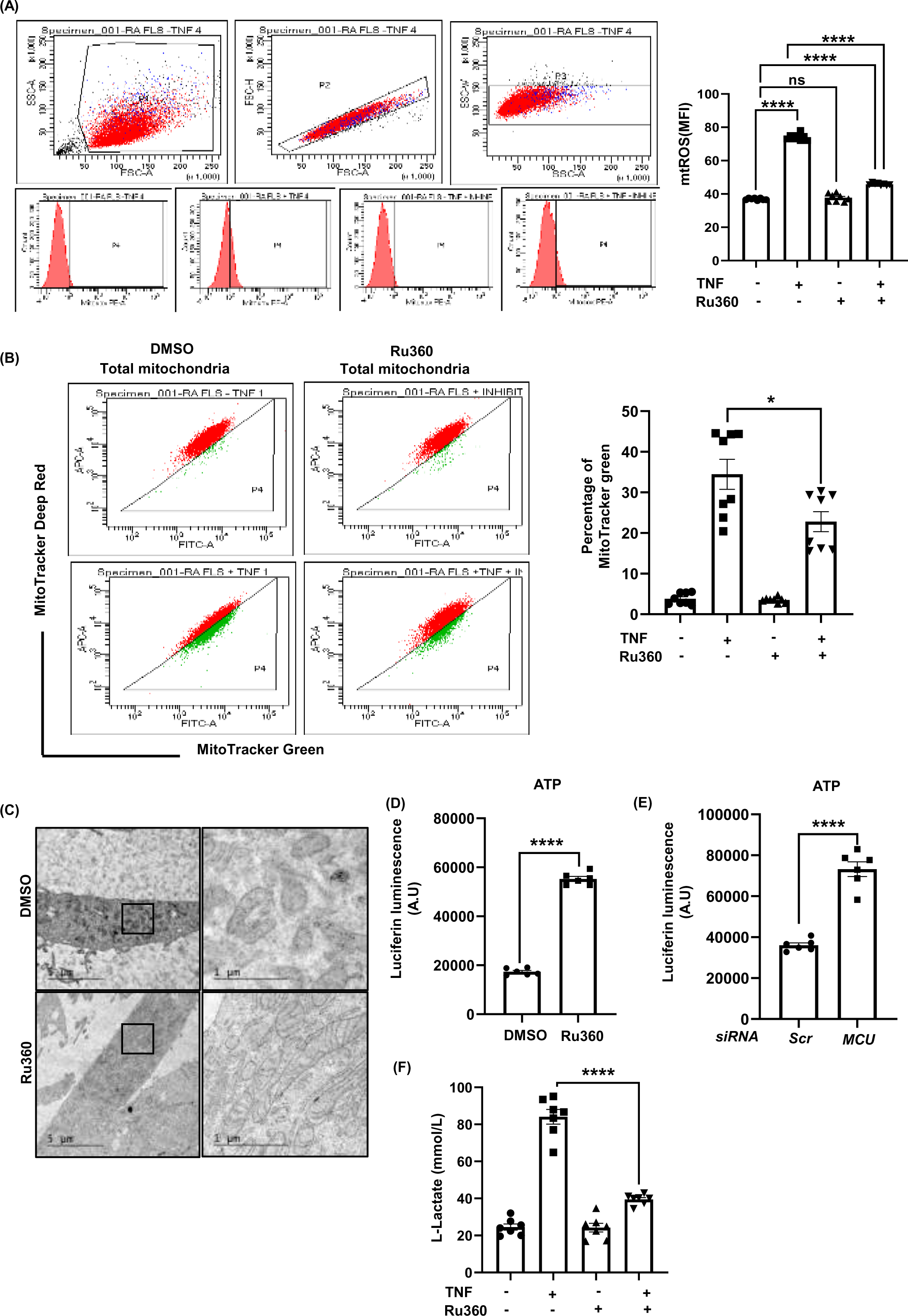
Effect of MCU inhibition on mitochondrial function: (**A**) Mitochondrial ROS production by FACS in RAFLS, isolated from rheumatoid arthritis patients (n=8). Cells were incubated with 5 μM Mitosox RED for 20 min at 37°. After trypsinization and centrifugation at 4°C the cells were analyzed for Mitosox signal in a flow cytometer. Representative FACS image of RAFLS treated with or without TNF (20ng/ml) and Ru360 (3uM). Bar graph indicates median value ± SD of Mitosox positive cells in RAFLS cells. **(B)** Transmission electron microscopy of RAFLS cells treated with and without Ru360, showing the ultrastructure of the mitochondria. **(C)** RAFLS cells were treated with or without TNF (20ng/ml) and Ru360 (3uM) for 24 hours, and stained with MitoTracker Deep Red (for healthy mitochondria) and MitoTracker Green (for total mitochondria) for 30 minutes to assess mitochondrial dysfunction by flow cytometry. Representative and summarized flow cytometry with the mean fluorescence intensity (MFI) values are shown (pooled data from n = 8 patients are shown). **(D)** Determination of L-Lactate concentration by human RA-FLS (n=7) after inhibition of MCU with Ru360. **(E, F)** The bar graph illustrates the ATP producing capacity by human RA-FLS (n=6) after being treated with Ru360 (3uM) and *siMCU* (100nM). The y-axis depicts the luciferase bioluminescence (ATP) in arbitrary units. Statistical significance was determined using t-test. *\*p < 0.05, **p < 0.01, ***p < 0.001, ****p < 0.0001*.

We next turned to the question how an inhibitor improves the mitochondrial function. It is widely known that ultrastructure of mitochondria highly disrupts in the diseases related to the mitochondria. Moreover, mitochondrial cristae morphology reflects the superoxide formation, metabolism, homeostasis and pathology **(Plecita-Hlavata and Jezek, 2016)**. To complete the picture, human RA-FLS cells were treated with Ru360 and compared to those without inhibitor and proceeded for electron microscopy to determine the ultrastructure. Interestingly, our data showed a remarkable improvement in the ultrastructure of the cristae after MCU inhibition (**figure 5C**).

It has long been established that an improved cristae structure indicates better mitochondrial health and function, which is crucial because cristae integrity is essential for ATP production **(Venkatraman et al., 2023)**. To accomplish this, we measured the amount of ATP generated by the cells with and without Ru360 treatment. As expected, we observed the increased ATP production in the inhibitor group as compared to those without inhibitor in human RA-FLS cells (**Figure 5D-E**). However, we still did not know whether the ATP produced was from the glycolytic pathway or oxidative phosphorylation. To verify this, we investigated glycolysis by measuring lactate production **(Young et al., 2013)**. Noteworthy, we observed that, upon Ru360, **L-lactate** production was reduced in human RA-FLS cells compared to without inhibitor (**Figure 5F**). Taken together, our data indicates a metabolic shift from glycolysis to oxidative phosphorylation, suggesting an improved mitochondrial function in the presence of MCU inhibition.

As we are looking for the effect of MCU inhibition on mitochondria, we decided to focus further on the metabolic intermediates. Hence, for this reason we screened the metabolomics of the human RA-FLS under four conditions: +TNF, –TNF, +TNF –Ru360, and +TNF +Ru360. The image focuses on the top 25 enriched metabolite sets, highlighting their potential involvement in the biological process under investigation. The significant enrichment in pathways like fructose and mannose degradation, glycolysis, gluconeogenesis, and lactose degradation indicates a shift towards increased glucose metabolism. This is crucial for cellular energy production and proliferation. Glycolysis, in particular, provides energy and intermediates for biosynthesis, supporting rapid cell division and growth **(Vander Heiden et al., 2009)**. Enhanced glycolysis and gluconeogenesis can drive cell proliferation by supplying ATP and metabolic intermediates needed for nucleotide and lipid biosynthesis **(Liberti and Locasale, 2016)**. Enrichment in sphingolipid and glycerolipid metabolism suggests alterations in membrane composition and signaling pathways. Sphingolipids are involved in regulating cell growth, apoptosis, and migration **(Ogretmen, 2018)**. Glycerolipids play a role in membrane formation and energy storage, impacting cell migration and invasion **(Fu et al., 2021)**. Changes in lipid metabolism can influence membrane fluidity and signaling pathways, thus affecting cell motility and invasiveness. The significant involvement of arginine and proline metabolism can affect cell proliferation and migration. Arginine is a precursor for nitric oxide (NO), which influences cell proliferation and migration through various signaling pathways (Bogdan, 2001). Proline metabolism is linked to cellular stress responses and collagen synthesis, affecting cell adhesion and migration **(Phang et al., 2010)**. Overall, the **figure 6**, illustrates that TNF treatment enhances multiple metabolic pathways, including glycolysis, the pentose phosphate pathway, the Krebs cycle, purine metabolism, and pyrimidine metabolism in RA-FLS. Inhibition of MCU reverses these metabolic changes, highlighting the role of MCU in promoting metabolic alterations associated with inflammation and possibly disease progression in rheumatoid arthritis.

**Figure 6:**
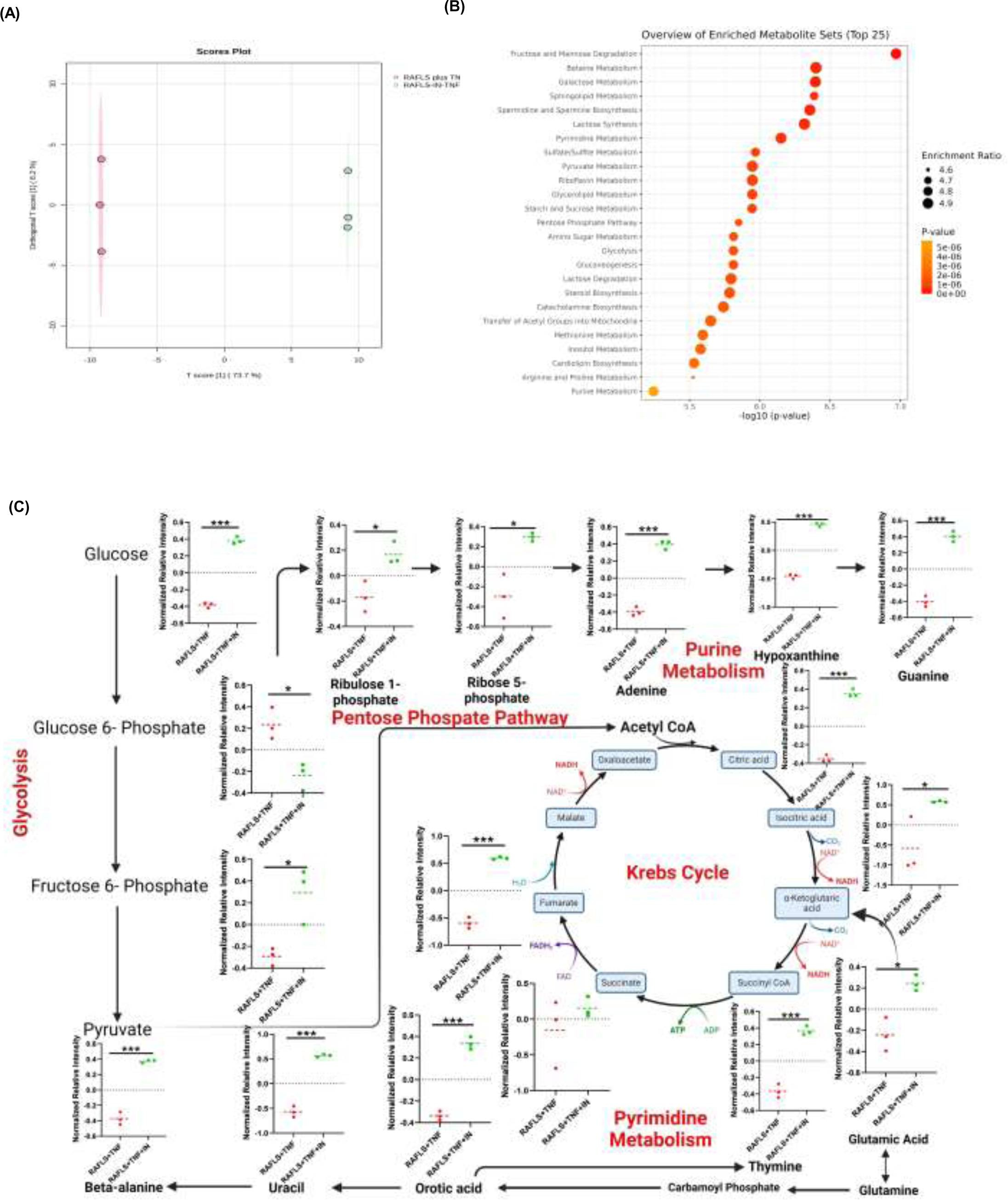
Metabolite profiling of RA-FLS cells under TNF stimulation and MCU inhibition: (**A**) Heatmap of metabolite levels in RAFLS-treated cells exposed to TNF treated with or without TNF (RAFLS-IN-TNF plus). Red indicates increased levels; blue indicates decreased levels relative to the mean. **(B)** Scatter plot showing Comparison of metabolite scores between RAFLS plus TNF and RAFLS-IN-TNF plus conditions. Each dot represents a metabolite. Green dots indicate metabolites significantly altered by TNF stimulation. **(C)** Volcano plot of metabolite fold change (FC) and p-value. Each dot represents a metabolite. The x-axis shows the log2(FC) between with or without inhibitor in presence of TNF. The y-axis shows the –log10(p-value) of the differential abundance test. Metabolites with a p-value < 0.05 and |log2(FC)| > 1 are considered significantly altered and are highlighted. Red dots indicate upregulated metabolites, while blue dots indicate downregulated metabolites. (**D)** VIP scores of metabolites differentiating RAFLS plus TNF and RAFLS-IN-TNF plus conditions. Metabolites are ranked by VIP score, a measure of their importance in discriminating between the two conditions. The heatmap on the right display metabolite levels (z-scores) in RAFLS plus TNF (red) and RAFLS-IN-TNF plus (blue) conditions. Higher VIP scores indicate greater contribution of the metabolite to the differentiation. (**E)** Overview of enriched metabolite sets. The dot plot illustrates the top 25 enriched metabolite sets based on enrichment ratio and p-value. Each dot represents a metabolite set, with size and color indicating enrichment ratio and p-value, respectively. Larger dot size and warmer colors denote higher enrichment ratios and lower p-values, suggesting greater significance.

### Miro-1 interacts with MCU to modulate the mitochondria localization

Recent studies revealed that miro-1 is the protein which takes the mitochondria to the leading edge and the interaction between miro-1 and MCU is very well characterized **(Niescier et al., 2018)**. So, to verify whether the effect on migration is MCU-dependent or not. We first overexpress the cells with MCU plasmid followed by the pull-down to validate the interaction between both. Our data clearly shows the interaction between MCU and miro-1 in **figure 7(A)**.

**Figure 7:**
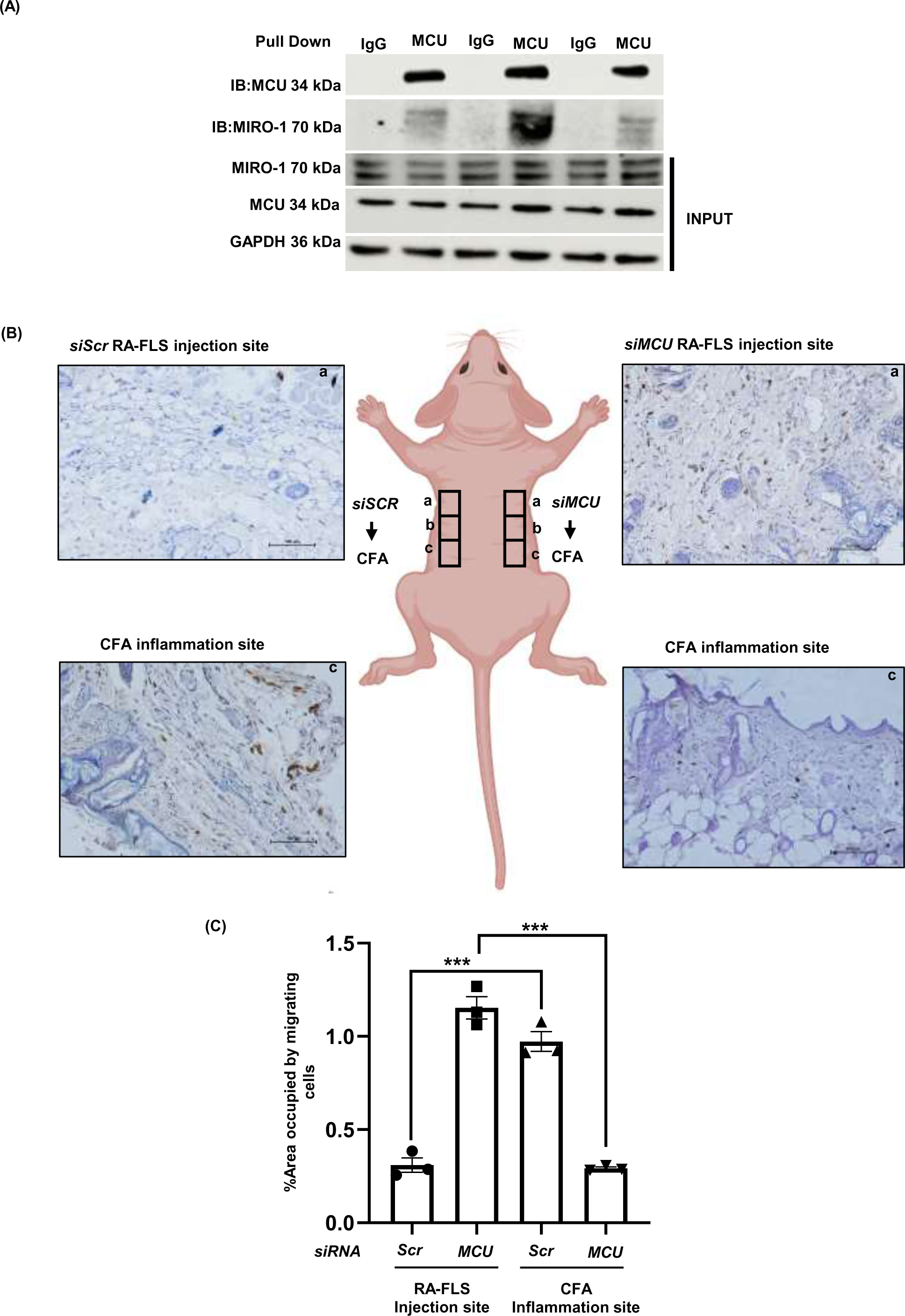
MCU interacts with Miro1 and MCU inhibition suppresses FLS migration in vivo. (**A**) Western blot analysis demonstrating MCU-Miro1 interaction. Whole cell lysates were immunoprecipitated (IP) with MCU antibody followed by Western blot analysis with Miro1 antibody. GAPDH served as a loading control. The presence of Miro1 in the MCU immunoprecipitated indicates an interaction between the two proteins. **(B)** A schematic xenograft model of migration assay. In athymic nude mice, skin inflammation was induced by subcutaneously injecting 120 µg of complete Freund’s adjuvant (CFA) into the site ‘c’. a day subsequent to CFA injection, human RA-FLS (5×10^5^) *siScr* and human RA-FLS (5×10^5^) *siMCU* cells were injected at site ‘a’ at a distance 1.5cm away from site ‘c’. Following a period of 5 days post FLS implantation, skin specimens were collected from specific locations ‘a’ and ‘c’ from both the sides. Increased migration of RA-FLS (DAB stained) was detected in the skin obtained from site ‘c’ of left side and less migration of *siMCU* RA-FLS from site (a) of right side, as determined by immuno-histochemical staining using anti-HLA class I antibody. DAB-stained cells were counted using ImageJ. **(C)** A bar graph below the schematic diagram quantifies the percentage of area occupied by migrating cells at each site. Comparisons are made between the *siSCR* RAFLS injection site, *siMCU* RAFLS injection site, and CFA inflammation sites. Values are the mean ± SEM of 3 mice. *\*, p < 0.0001* versus control *siRNA*-transfected FLS.

Miro1 and MCU are known to work together to regulate the migration of mitochondria to the leading edge of cells **(Niescier et al., 2018)**. We inhibited Miro1 using a Miro1 reducer and then evaluated whether the cells migrated. This experiment was designed to determine whether the disruption of the MCU-Miro1 interaction would alter the migration process in RA-FLS cells, providing insights into the connection between Miro1, MCU, and cellular movement in the context of rheumatoid arthritis. Upon inhibiting Miro-1, we observed a significant reduction in the migration of RA-FLS cells (data not shown). This decrease suggests that Miro-1 is essential in promoting cell movement.

### MCU silencing hampers the metastasis in athymic nude mice

The *in vitro* data on migration and invasiveness were further validated by an *in vivo* experiment. *siScr* and *siMCU* silenced RA-FLS cells were injected subcutaneously into the SCID mice on the left and right side respectively. One day before the cells injection, Complete Freund’s Adjuvant (CFA) was induced 1.5cm away from the RA-FLS injection site. Five days after injecting the RA-FLS cells, skin tissue samples were taken from both the area where the RA-FLS were injected and the site where CFA was administered. A cluster of human RA-FLS cells (HLA-DR positive) was found more often in the skin where CFA was injected on the left side when using control *siRNA* compared to the right side where MCU *siRNA* was used, suggesting that MCU inhibition reduces FLS migration toward the CFA-injected site with dermal. Overall, these data suggest that MCU siRNA suppresses FLS migration *in vivo* **figure 7(B),(C).**

## Discussion

Despite significant advancements in medical science, rheumatoid arthritis remains an incurable due to its complex and multifaceted origins. As a variety of cells participate in establishment and perpetuation of RA. Notably, FLS are the major contributor to the pathogenesis of the rheumatoid arthritis (RA). The joint pathology of Rheumatoid Arthritis (RA) is characterized by an overgrowth of FLS that invades and damages nearby cartilage and bones. RA –FLS contributes to the infiltration of inflammatory immune cells in the synovial space and drives inflammation. RA-FLS are known to produce chemokines, proinflammatory cytokines, and MMPs, the enzymes that break down extracellular matrix tissue and helps in the migration and invasion. We believe that a deeper understanding of RA-FLS behavior could identify new therapeutic targets for RA. This increased invasiveness is linked to worse disease progression and joint damage. A research showed that a protein complex called MCU helps to control migration of tumor cells by reducing mitochondrial calcium uptake **(Paupe and Prudent, 2018)**. Recently, how calcium ion uptake ([Ca2+]m) contributes to tumor growth and the spread of cancer (metastasis) has been extensively studied. Studies have identified MCU as the primary selective channel responsible for transporting calcium ions cytosol to matrix of mitochondria (**Baughman et al., 2011**). This discovery has led to investigations into the role of calcium ion uptake in various diseases, including cancer. In this context one research has demonstrated a link between MCU expression and cancer metastasis (**Tosatto et al., 2016)**. However, the exact way MCU worked was unclear. Within this framework, our study sheds new light on the pathogenesis of rheumatoid arthritis (RA) by elucidating the pivotal role of the mitochondrial calcium uniporter (MCU), the major contributor of mitochondrial calcium influx, in regulating the invasiveness and migration of rheumatoid arthritis fibroblast-like synoviocytes (RA-FLS). By inhibiting the MCU, we observed a significant reduction in the migration and invasion capabilities of RA-FLS, suggesting that calcium uptake through the MCU is critical for these pathological processes. Our study found a new mechanism by which MCU regulates RA-FLS behavior. We discovered that MCU inhibition reduces specific proteins such as RhoA and Rac1, and actin filaments polymerization which are involved in cell movement and attachment. Rac1 has been identified as a key regulator in cancer development (**Chan et al., 2005**) as well as influencing the invasiveness of FLS (**Chan et al., 2007**). This protein plays a crucial role in restructuring the FLS cytoskeleton, enabling the cell to spread out and form cellular extensions called lamellipodia (**Chan et al., 2007**). By doing so, MCU inhibition reduces the ability of RA-FLS to adhere to and invade surrounding tissue. Moreover, Increasing the levels of MCU in MCF-7 breast cancer cells led to increased migratory and invasive capabilities *in vitro* and lung metastasis in an *in vivo* mouse model (**Yu et al., 2017**). To support the hypothesis we have reversed the study and observed the similar results, the overexpression of MCU leads to an overload of calcium that supports more invasion of RA-FLS. Interestingly, mitochondria play a central role in the migration of any migrating cell. Mitochondria, which are the organelles in charge of synthesizing ATP aerobically, create a tubular network that merges with different structures within the cell. As different regions of the cell experience varying energy demands, mitochondria are fragmented by Drp1 into more mobile structures that are relocated to where they are most needed. The process of mitochondrial division is vital for preserving the normal physiological activities of cells. However, cancer cells exhibit atypical bioenergetics, often obtaining a significant portion of their energy from aerobic glycolysis (**Jose et al., 2011)**, even when oxygen is available. In a study, breast tissue samples has been examined and found that the amount of Drp1 protein was slightly higher in non-invasive breast cancer compared to normal tissue. However, there was a significant increase in Drp1 protein in invasive breast cancer and cancer that had spread to lymph nodes. These findings suggest that an increase in Drp1 protein might be an early sign of breast cancer becoming invasive (**Zhao et al., 2013)**. Interestingly, during migration mitochondria becomes small due to higher expression of fission genes Drp-1 (**Kim et al., 2015**) and lower expression of fusion genes including Mfn-1. **(Zhao et al., 2013**). When comparing the fission and fusion markers in FLS cells isolated from CIA mice, we observed similar results. To unleash the mechanism behind the migration of RA-FLS cells, we focused on the genes responsible for the migration of FLS cells post MCU inhibition. A variety of genes were upregulated and downregulated after MCU inhibition. Interestingly, our transcriptome analysis highlighted marked downregulation of actin cytoskeleton and focal adhesion genes, which are crucial for cell motility, adhesion, and invasion. To complete the picture, we measured the Rac1 activity after MCU inhibition and observed a significant reduction in post-Ru360 treatment, further confirming the reduced cell migration. Not only that, mitochondria are the primary source of reactive oxygen species (ROS) also within cells. ROS can act as signaling molecules, stimulating cell growth and migration (P**elicano et** al., 2009),(**Luanpitpong et al., 2010**). However, the process by which ROS enhances cell motility is not yet fully understood. Consistent with this, we hypothesized that inhibiting calcium uptake would reduce mitochondrial ROS. Treating human RA-FLS cells with Ru360 under TNF conditions resulted in a significant reduction in mitochondrial ROS production, indicating Ru360 as a potent inhibitor for reducing mitochondrial ROS in inflammation.

The production and accumulation of ROS contribute to the damage of mitochondrial DNA, proteins, and lipids. This damage impairs the mitochondria’s ability to function properly, leading to decreased efficiency in energy production and further increased ROS generation. As this cycle continues, the accumulating damage exacerbates mitochondrial dysfunction (**Dan Dunn et al., 2015**) lead to impaired mitochondrial cristae structure. Cristae are formed through the invagination of the inner mitochondrial membrane, creating a large surface area necessary for ATP production. The structural integrity and organization of cristae are vital for efficient mitochondrial function. A study has revealed that calcium overload significantly disrupt mitochondrial ultrastructure. Under normal conditions, mitochondria have a tightly packed, complex internal membrane system, essential for energy production and maintain an intact and dense cristae network. However, when mitochondria are overwhelmed by calcium the ultrastructure get disturbed and causes substantial remodeling of the inner membrane, disrupting the cristae formation and organization, leading to a decline in the cell’s energy generation capacity. (**Strubbe-Rivera et al., 2021**). Taking this into account, we have analyzed the mitochondrial ultrastructure of human RA-FLS. We observed that some mitochondria had disturbed cristae. However, when the RA-FLS were treated with Ru360, the structure showed some improvement. Additionally, mitochondria were comparatively more fragmented without Ru360 treatment, while fused mitochondria were comparatively more common with the treatment. Subsequently, we measured mitochondrial function by examining glycolysis. This is important because tumor cells, unlike normal cells, rely more heavily on glycolysis to fulfill their ATP production requirements **(Shiratori et al., 2019).** Surprisingly, lactate production was reduced following Ru360 treatment, suggesting a shift in metabolism towards oxidative phosphorylation. Consequently, we measured ATP production and observed a marked increase in ATP levels after treating the cells with Ru360. Glycolytic suppression significantly alters the intracellular metabolic profile of various cancer cell lines within mitochondria (**Shiratori et al., 2019**).

The presented metabolomic analysis reveals a profound impact of TNF on the metabolic landscape of RA-FLS cells, with a pronounced upregulation of glycolytic and lipogenic pathways. These metabolic shifts are consistent with the well-established Warburg effect, a hallmark of cancer and inflammatory diseases**(Burns and Manda, 2017).** The observed enrichment in pathways related to glucose metabolism, nucleotide biosynthesis, and lipid synthesis underscore the heightened metabolic demands of proliferating and activated cells. Importantly, the inhibition of MCU, a key regulator of mitochondrial Ca^2+^ influx, effectively reversed these metabolic alterations. This finding strongly implicates MCU as a critical mediator of metabolic reprogramming in RA-FLS. By attenuating MCU activity, we observed a suppression of glycolysis, lipid synthesis, and related pathways, suggesting a potential therapeutic strategy to target aberrant metabolism in rheumatoid arthritis.The observed changes in sphingolipid and glycerolipid metabolism highlight the complex interplay between energy metabolism and cellular signaling. Altered lipid composition can profoundly impact membrane fluidity, receptor signaling, and inflammatory responses. The involvement of arginine and proline metabolism further emphasizes the intricate connections between metabolic pathways and cellular functions. In conclusion, our data provide compelling evidence for the central role of MCU in driving metabolic reprogramming in RA-FLS cells under inflammatory conditions. A recent study strongly suggests that MCU is crucial for controlling both the entry of calcium ions into mitochondria and the movement of mitochondria along nerve cell fibers (axons). The researchers found that MCU interacts with a protein called Miro1, located on the outer membrane of mitochondria (**Fransson et al., 2006**), through an N-terminal mitochondrial targeting domain. Surprisingly, this N-terminal domain of MCU isn’t solely responsible for its positioning within the cell. The study indicates that the interaction between MCU and Miro1 is vital for the efficient movement of mitochondria in axons but not for the intake of calcium into mitochondrial matrix **(Niescier et al., 2018**). Considering this, we first confirmed the interaction between Miro-1 and MCU through co-immunoprecipitation. Subsequently, we aimed to inhibit Miro-1 using a Miro-1 reducer and observed a reduction in cell migration of human RA-FLS cells. This suggests an MCU-independent function of Miro-1 in regulating cell migration. Thus, Miro-1 is a protein that can also potentially regulate the migration of human RA-FLS cells independently of MCU.

Our findings demonstrate a critical role for MCU in regulating RA-FLS migration and invasion *in vitro*. By inhibiting MCU expression using *siRNA*, we observed a significant reduction in both migratory and invasive capacities of RA-FLS cells. Furthermore, *in vivo* experiments confirmed these results, showing that MCU-silenced RA-FLS cells exhibited decreased migration toward a site of inflammation compared to control cells. These data collectively suggest that MCU is a key regulator of RA-FLS migration and may represent a promising therapeutic target for rheumatoid arthritis. By inhibiting MCU, we propose a potential strategy to impede the progression of rheumatoid arthritis by reducing the infiltration and accumulation of RA-FLS in inflamed tissues.

Suggesting, targeting the MCU could offer a novel therapeutic strategy for RA. Current RA treatments, including disease-modifying antirheumatic drugs (DMARDs) and biologics, primarily focus on suppressing inflammation and modulating immune responses. However, these treatments do not directly address the invasive properties of RA-FLS. Our findings suggest that MCU inhibitors could complement existing therapies by specifically targeting the cellular mechanisms that drive joint destruction. Further research is needed to explore the therapeutic potential of MCU inhibitors in clinical settings. Our study opens several avenues for future research.

## Conclusion

In conclusion, our study identifies the mitochondrial calcium uniporter as a key regulator of RA-FLS migration and invasion, thus playing a significant role in the pathogenesis of rheumatoid arthritis. By inhibiting the MCU, we can disrupt the invasive behavior of RA-FLS, suggesting a promising new target for therapeutic intervention. These findings enhance our understanding of RA pathogenesis and pave the way for the development of novel treatments aimed at mitigating joint destruction and improving patient outcomes in rheumatoid arthritis.

## Abbreviations

RA: rheumatoid arthritis
RA-FLS: rheumatoid arthritis fibroblast-like synoviocytes
MCU: mitochondrial calcium uniporter
RR: Ruthenium red
Ru360: Ruthenium 360
TEM: transmission electron microscopy
DMARDs: Disease-Modifying Anti-Rheumatic Drugs.
JAK: Janus kinase
MTX: Methotrexate
MMP: Matrix metalloproteinases
TNF: tumour necrosis factor
HIP1: Huntingtin-interacting protein 1
TRPV2: transient receptor potential vanilloid subfamily, type 2 channel

## Acknowledgments

The work is supported by financial grants from the ICMR Ad-hoc (Ortho) 2020-NCD-1; CSIR HCP-0047. We gratefully received technical help from Mr. Anil Verma for microscopy and Deviprasad Panda for flow cytometry studies and Sophisticated Analytical Instrument Facility and Research Division, CSIR – Central Drug Research Institute. CSIR THUNDER (BSC0102) and MOES (GAP0118) Intravital Facility and Laboratory Animal Facility of CSIR – Central Drug Research Institute are acknowledged.PL, MR and KS are funded by CSIR and SG, NK and MST are funded by UGC. VA is funded by DBT Wellcome Trust India Alliance (IA/E/21/1/506319).This manuscript has a CSIR – Central Drug Research Institute communication number: **127/2024/AL**

## Specific contribution of each author

PL, KS, SG, SK, MR, NCS, MST have performed the experiments, analyzed data and drafted the paper. MKB, JK, AA, RG, VA, AS, KM, AL supervised the experiments and analyzed the data, wrote the paper.

## References

1. Baughman, J.M., F. Perocchi, H.S. Girgis, M. Plovanich, C.A. Belcher-Timme, Y. Sancak, X.R. Bao, L. Strittmatter, O. Goldberger, R.L. Bogorad, V. Koteliansky, and V.K. Mootha. 2011. Integrative genomics identifies MCU as an essential component of the mitochondrial calcium uniporter. Nature. 476:341–345.

2. Bian, Y., Z. Xiang, Y. Wang, Q. Ren, G. Chen, B. Xiang, J. Wang, C. Zhang, S. Pei, S. Guo, and L. Xiao. 2023. Immunomodulatory roles of metalloproteinases in rheumatoid arthritis. Front Pharmacol. 14:1285455.

3. Bogdan, C. 2001. Nitric oxide and the immune response. Nat Immunol. 2:907–916.

4. Bottini, N., and G.S. Firestein. 2013. Duality of fibroblast-like synoviocytes in RA: passive responders and imprinted aggressors. Nat Rev Rheumatol. 9:24–33.

5. Burns, J.S., and G. Manda. 2017. Metabolic Pathways of the Warburg Effect in Health and Disease: Perspectives of Choice, Chain or Chance. Int J Mol Sci. 18.

6. Bustamante, M.F., R. Garcia-Carbonell, K.D. Whisenant, and M. Guma. 2017. Fibroblast-like synoviocyte metabolism in the pathogenesis of rheumatoid arthritis. Arthritis Res Ther. 19:110.

7. Chan, A., M. Akhtar, M. Brenner, Y. Zheng, P.S. Gulko, and M. Symons. 2007. The GTPase Rac regulates the proliferation and invasion of fibroblast-like synoviocytes from rheumatoid arthritis patients. Mol Med. 13:297–304.

8. Chan, A.Y., S.J. Coniglio, Y.Y. Chuang, D. Michaelson, U.G. Knaus, M.R. Philips, and M. Symons. 2005. Roles of the Rac1 and Rac3 GTPases in human tumor cell invasion. Oncogene. 24:7821–7829.

9. Chen, W., H. Zhao, and Y. Li. 2023. Mitochondrial dynamics in health and disease: mechanisms and potential targets. Signal Transduct Target Ther. 8:333.

10. Dan Dunn, J., L.A. Alvarez, X. Zhang, and T. Soldati. 2015. Reactive oxygen species and mitochondria: A nexus of cellular homeostasis. Redox Biol. 6:472–485.

11. Dang, J., S. Zhu, and J. Wang. 2019. A protocol for humanized synovitis mice model. Am J Clin Exp Immunol. 8:47–52.

12. De Stefani, D., M. Patron, and R. Rizzuto. 2015. Structure and function of the mitochondrial calcium uniporter complex. Biochim Biophys Acta. 1853:2006–2011.

13. Dubinsky, J.M., and Y. Levi. 1998. Calcium-induced activation of the mitochondrial permeability transition in hippocampal neurons. J Neurosci Res. 53:728–741.

14. Fransson, S., A. Ruusala, and P. Aspenstrom. 2006. The atypical Rho GTPases Miro-1 and Miro-2 have essential roles in mitochondrial trafficking. Biochem Biophys Res Commun. 344:500–510.

15. Fu, Y., T. Zou, X. Shen, P.J. Nelson, J. Li, C. Wu, J. Yang, Y. Zheng, C. Bruns, Y. Zhao, L. Qin, and Q. Dong. 2021. Lipid metabolism in cancer progression and therapeutic strategies. MedComm (2020). 2:27–59.

16. Guma, M., S. Tiziani, and G.S. Firestein. 2016. Metabolomics in rheumatic diseases: desperately seeking biomarkers. Nat Rev Rheumatol. 12:269–281.

17. Hall, A., and C.D. Nobes. 2000. Rho GTPases: molecular switches that control the organization and dynamics of the actin cytoskeleton. Philos Trans R Soc Lond B Biol Sci. 355:965–970.

18. Haroon, N., A. Aggarwal, A. Lawrence, V. Agarwal, and R. Misra. 2007. Impact of rheumatoid arthritis on quality of life. Mod Rheumatol. 17:290–295.

19. Itoh, R.E., E. Kiyokawa, K. Aoki, T. Nishioka, T. Akiyama, and M. Matsuda. 2008. Phosphorylation and activation of the Rac1 and Cdc42 GEF Asef in A431 cells stimulated by EGF. J Cell Sci. 121:2635–2642.

20. Jose, C., N. Bellance, and R. Rossignol. 2011. Choosing between glycolysis and oxidative phosphorylation: a tumor’s dilemma? Biochim Biophys Acta. 1807:552–561.

21. Kim, H.J., M.R. Shaker, B. Cho, H.M. Cho, H. Kim, J.Y. Kim, and W. Sun. 2015. Dynamin-related protein 1 controls the migration and neuronal differentiation of subventricular zone-derived neural progenitor cells. Sci Rep. 5:15962.

22. Koedderitzsch, K., E. Zezina, L. Li, M. Herrmann, and N. Biesemann. 2021. TNF induces glycolytic shift in fibroblast like synoviocytes via GLUT1 and HIF1A. Sci Rep. 11:19385.

23. Koval, O.M., E.K. Nguyen, V. Santhana, T.P. Fidler, S.C. Sebag, T.P. Rasmussen, D.J. Mittauer, S. Strack, P.C. Goswami, E.D. Abel, and I.M. Grumbach. 2019. Loss of MCU prevents mitochondrial fusion in G(1)-S phase and blocks cell cycle progression and proliferation. Sci Signal. 12.

24. Kristal, B.S., and J.M. Dubinsky. 1997. Mitochondrial permeability transition in the central nervous system: induction by calcium cycling-dependent and –independent pathways. J Neurochem. 69:524–538.

25. Laragione, T., M. Brenner, A. Lahiri, E. Gao, C. Harris, and P.S. Gulko. 2018. Huntingtin-interacting protein 1 (HIP1) regulates arthritis severity and synovial fibroblast invasiveness by altering PDGFR and Rac1 signalling. Ann Rheum Dis. 77:1627–1635.

26. Laragione, T., M. Brenner, A. Mello, M. Symons, and P.S. Gulko. 2008. The arthritis severity locus Cia5d is a novel genetic regulator of the invasive properties of synovial fibroblasts. Arthritis Rheum. 58:2296–2306.

27. Laragione, T., C. Harris, and P.S. Gulko. 2019. TRPV2 suppresses Rac1 and RhoA activation and invasion in rheumatoid arthritis fibroblast-like synoviocytes. Int Immunopharmacol. 70:268–273.

28. Liberti, M.V., and J.W. Locasale. 2016. The Warburg Effect: How Does it Benefit Cancer Cells? Trends Biochem Sci. 41:211–218.

29. Luanpitpong, S., S.J. Talbott, Y. Rojanasakul, U. Nimmannit, V. Pongrakhananon, L. Wang, and P. Chanvorachote. 2010. Regulation of lung cancer cell migration and invasion by reactive oxygen species and caveolin-1. J Biol Chem. 285:38832–38840.

30. Ma, N., E. Xu, Q. Luo, and G. Song. 2023. Rac1: A Regulator of Cell Migration and a Potential Target for Cancer Therapy. Molecules. 28.

31. Mousavi, M.J., J. Karami, S. Aslani, M.N. Tahmasebi, A.S. Vaziri, A. Jamshidi, E. Farhadi, and M. Mahmoudi. 2021. Transformation of fibroblast-like synoviocytes in rheumatoid arthritis; from a friend to foe. Auto Immun Highlights. 12:3.

32. Muller-Ladner, U., R.E. Gay, and S. Gay. 2000. Activation of synoviocytes. Curr Opin Rheumatol. 12:186–194.

33. Niescier, R.F., K. Hong, D. Park, and K.T. Min. 2018. MCU Interacts with Miro1 to Modulate Mitochondrial Functions in Neurons. J Neurosci. 38:4666–4677.

34. Ogretmen, B. 2018. Sphingolipid metabolism in cancer signalling and therapy. Nat Rev Cancer. 18:33–50.

35. Papalazarou, V., and L.M. Machesky. 2021. The cell pushes back: The Arp2/3 complex is a key orchestrator of cellular responses to environmental forces. Curr Opin Cell Biol. 68:37–44.

36. Paupe, V., and J. Prudent. 2018. New insights into the role of mitochondrial calcium homeostasis in cell migration. Biochem Biophys Res Commun. 500:75–86.

37. Pelicano, H., W. Lu, Y. Zhou, W. Zhang, Z. Chen, Y. Hu, and P. Huang. 2009. Mitochondrial dysfunction and reactive oxygen species imbalance promote breast cancer cell motility through a CXCL14-mediated mechanism. Cancer Res. 69:2375–2383.

38. Phang, J.M., W. Liu, and O. Zabirnyk. 2010. Proline metabolism and microenvironmental stress. Annu Rev Nutr. 30:441–463.

39. Plecita-Hlavata, L., and P. Jezek. 2016. Integration of superoxide formation and cristae morphology for mitochondrial redox signaling. Int J Biochem Cell Biol. 80:31–50.

40. Porporato, P.E., V.L. Payen, J. Perez-Escuredo, C.J. De Saedeleer, P. Danhier, T. Copetti, S. Dhup, M. Tardy, T. Vazeille, C. Bouzin, O. Feron, C. Michiels, B. Gallez, and P. Sonveaux. 2014. A mitochondrial switch promotes tumor metastasis. Cell Rep. 8:754–766.

41. Price, L.S., M. Langeslag, J.P. ten Klooster, P.L. Hordijk, K. Jalink, and J.G. Collard. 2003. Calcium signaling regulates translocation and activation of Rac. J Biol Chem. 278:39413–39421.

42. Promila, L., A. Joshi, S. Khan, A. Aggarwal, and A. Lahiri. 2023. Role of mitochondrial dysfunction in the pathogenesis of rheumatoid arthritis: Looking closely at fibroblast-like synoviocytes. Mitochondrion. 73:62–71.

43. Shiratori, R., K. Furuichi, M. Yamaguchi, N. Miyazaki, H. Aoki, H. Chibana, K. Ito, and S. Aoki. 2019. Glycolytic suppression dramatically changes the intracellular metabolic profile of multiple cancer cell lines in a mitochondrial metabolism-dependent manner. Sci Rep. 9:18699.

44. Strubbe-Rivera, J.O., J. Chen, B.A. West, K.N. Parent, G.W. Wei, and J.N. Bazil. 2021. Modeling the Effects of Calcium Overload on Mitochondrial Ultrastructural Remodeling. Appl Sci (Basel). 11.

45. Tang, S., X. Wang, Q. Shen, X. Yang, C. Yu, C. Cai, G. Cai, X. Meng, and F. Zou. 2015. Mitochondrial Ca(2)(+) uniporter is critical for store-operated Ca(2)(+) entry-dependent breast cancer cell migration. Biochem Biophys Res Commun. 458:186–193.

46. Tosatto, A., R. Sommaggio, C. Kummerow, R.B. Bentham, T.S. Blacker, T. Berecz, M.R. Duchen, A. Rosato, I. Bogeski, G. Szabadkai, R. Rizzuto, and C. Mammucari. 2016. The mitochondrial calcium uniporter regulates breast cancer progression via HIF-1alpha. EMBO Mol Med. 8:569–585.

47. Vander Heiden, M.G., L.C. Cantley, and C.B. Thompson. 2009. Understanding the Warburg effect: the metabolic requirements of cell proliferation. Science. 324:1029–1033.

48. Venkatraman, K., C.T. Lee, G.C. Garcia, A. Mahapatra, D. Milshteyn, G. Perkins, K.Y. Kim, H.A. Pasolli, S. Phan, J. Lippincott-Schwartz, M.H. Ellisman, P. Rangamani, and I. Budin. 2023. Cristae formation is a mechanical buckling event controlled by the inner membrane lipidome. bioRxiv.

49. Wang, X., Z. Chen, X. Fan, W. Li, J. Qu, C. Dong, Z. Wang, Z. Ji, and Y. Li. 2020. Inhibition of DNM1L and mitochondrial fission attenuates inflammatory response in fibroblast-like synoviocytes of rheumatoid arthritis. J Cell Mol Med. 24:1516–1528.

50. Youle, R.J., and A.M. van der Bliek. 2012. Mitochondrial fission, fusion, and stress. Science. 337:1062–1065.

51. Young, S.P., S.R. Kapoor, M.R. Viant, J.J. Byrne, A. Filer, C.D. Buckley, G.D. Kitas, and K. Raza. 2013. The impact of inflammation on metabolomic profiles in patients with arthritis. Arthritis Rheum. 65:2015–2023.

52. Yu, C., Y. Wang, J. Peng, Q. Shen, M. Chen, W. Tang, X. Li, C. Cai, B. Wang, S. Cai, X. Meng, and F. Zou. 2017. Mitochondrial calcium uniporter as a target of microRNA-340 and promoter of metastasis via enhancing the Warburg effect. Oncotarget. 8:83831–83844.

53. Zeng, Y., M. Ren, Y. Li, Y. Liu, C. Chen, J. Su, B. Su, H. Xia, F. Liu, H. Jiang, H. Ling, X. Zeng, and Q. Su. 2020. Knockdown of RhoGDI2 represses human gastric cancer cell proliferation, invasion and drug resistance via the Rac1/Pak1/LIMK1 pathway. Cancer Lett. 492:136–146.

54. Zhao, J., J. Zhang, M. Yu, Y. Xie, Y. Huang, D.W. Wolff, P.W. Abel, and Y. Tu. 2013. Mitochondrial dynamics regulates migration and invasion of breast cancer cells. Oncogene. 32:4814–4824.

